# Characterizing the endopeptidase activity of *Candida albicans* Gpi8, a crucial subunit of the GPI transamidase

**DOI:** 10.64898/2026.04.07.717003

**Authors:** Isaac Cherian, Shailja Shefali, Dip Shikha Maurya, Faraz Mohd Khan, Sneha Sudha Komath

**Affiliations:** School of Life Sciences, Jawaharlal Nehru University, New Delhi-110067 India

**Keywords:** GPI transamidase, Gpi8/ PIG-K, cysteine protease, steady state kinetics, cell free assay, 7-amino-4-methylcoumarin (AMC)

## Abstract

GPI-anchored proteins are crucial cell surface proteins with diverse, organism-specific functions, in eukaryotes. They are produced when the GPI transamidase (GPIT), a five-subunit membrane-bound enzyme complex, attaches a pre-formed GPI anchor to the C-terminal end of nascent proteins on the lumenal face of the endoplasmic reticulum. This process requires the removal of a C-terminal signal sequence (SS) on the substrate protein by the action of an endopeptidase subunit of the GPIT, Gpi8/ PIG-K. Using an AMC-tagged peptide in a cell free (post-mitochondrial fraction) assay, this manuscript studies the steady state kinetics of enzymatic cleavage of the substrate by GPIT of the human pathogenic fungus, *C. albicans*. We show that Mn^+2^ enhances activity by improving substrate binding but plays no direct role in substrate cleavage *per se*. Molecular dynamics simulations suggest that the divalent cation binds at a site away from the active site but provides compactness and stability to Gpi8. It also enables a conformation in which a flexible loop (219-244 residues) in the vicinity of the catalytic pocket is able to interact with and position the scissile bond for cleavage by Cys202. Steady state kinetics also indicate that peptides of lengths 7-mer to 9-mer are better bound than 4-mer or 15-mer peptide substrates. A bulky residue at the site of cleavage reduces the catalytic activity of the GPIT. This is the first detailed steady state kinetics study on the endopeptidase activity of a GPIT from any organism.

## 1. Introduction

Glycosylphosphatidylinositol (GPI)-anchored proteins are cell surface proteins that play diverse roles in the eukaryotic cell. They could be enzymes, signal receptors, molecules involved in cell-cell interactions, or molecules involved in immune response [1]. The GPI anchor itself is a long glycolipid molecule synthesized in the endoplasmic reticulum (ER) of all eukaryotes by a sequential set of reactions, each catalyzed by a membrane-localized enzyme. Its biosynthesis begins at the cytoplasmic face of the ER with transfer of N-acetylglucosamine (GlcNAc) from UDP-GlcNAc to phosphatidylinositol (PI). GlcNAc-PI is then de-N-acetylated to glucosamine-PI (GlcN-PI), and flipped into the lumen, where the inositol is acylated and the carbohydrate chain extended by at least three Man residues, each of which is modified with a phosphoethanolamine (PEtN) residue [1]. In the fungal organisms, *C. albicans* and *S. cerevisiae,* a fourth Man is required to generate the GPI precursor, PEtN-6(Manα1-2)Manα1-2Manα1-6Manα1-4GlcNα1-6(acyl)myo-inositol-1-PO4-lipid [2,3]. Normally, the PEtN on Man-3 of these preformed glycolipids is finally amide-linked to the C-termini of proteins to be GPI-anchored after removal of their C-terminal GPI attachment signal sequences (SS). Both these jobs, are performed by a single enzyme, the GPI transamidase (GPIT), consisting of 5 subunits, Gpi8/ PIG-K, Gpi16/ PIG-T, Gaa1/ GPAA1, Gpi17/ PIG-S and Gab1/ PIG-U in *S. cerevisiae*/mammals. In *S. cerevisiae* the genes encoding the GPIT subunits are essential for viability, while in humans, defects in any of them result in disease [4]. Homologs of each of the subunits exist in *C. albicans*, a human pathogenic fungus that resides as a commensal in healthy individuals but turns opportunistic in immunocompromised conditions [4].

Once attached to the protein, the GPI anchor undergoes further modifications in the ER and the Golgi as it transits through the secretory pathway and reaches the cell surface where it is anchored in the external leaflet of the plasma membrane and/or the cell wall. Being linked to the cell wall implies that GPI-APs participate in cell wall integrity and any process that disturbs GPI biosynthesis also produces cell wall defects. Therefore, GPI biosynthetic defects and loss of GPI-APs in *C. albicans* produce cells that are more susceptible to cell wall damaging drugs/ agents while simultaneously exposing the underlying β-glucan components of the cell wall to attack and clearance by the host immune system [5,6]. Their ability to colonize and establish infection are also compromised since most of the host-recognition and virulence factors are GPI-APs [5]. Thus, enzymes of the GPI biosynthetic pathway are attractive antifungal drug targets and have been the focus of a new class of antifungals that are under development or in clinical trials in recent times [7]. This requires a detailed understanding of the catalytic mechanisms involved and the differences between the enzymes of the pathogen and host that can be exploited.

The fact that the GPIT is a multi-subunit membrane-bound enzyme makes attempts to purify or characterize it very challenging. Many previous studies depended on metabolic labeling and *in vitro* translation systems [8–10]. More recent efforts have been to obtain molecular details of the active site from cryo-EM structural data [11,12] and to develop more flexible enzyme assay strategies for recombinantly expressed soluble subunits/sub-complexes [13,14]. We previously showed that a heterozygous *C. albicans gpi8* strain accumulates CP intermediates and has reduced endopeptidase activity in a cell free assay, both of which could be reversed in a *GPI8* revertant strain [15]. Thus, the activity was specific to Gpi8. The cell free GPIT assay developed for this purpose used rough ER microsomes and a 7-residue fluorescently tagged peptide substrate, using which we found that the enzyme activity was sensitive to Cys- and His-modifying agents and was stimulated by the addition of Mn^+2^ and, to a lesser extent, by Ca^+2^ but not by Mg^+2^ or Zn^+2^.

Extending this study further, the present manuscript focuses on characterizing the enzymatic activity of *C. albicans* GPIT in post-mitochondrial subcellular fractions (PMF) and obtaining steady state kinetic parameters of the enzyme for different peptide substrates. From the results obtained, we conclude that the length of the peptide is an important criterion for substrate recognition by Gpi8. Substrates that are 7-mer and 9-mer in length have the most optimum K_M_ value, rather than a 4-mer or a 15-mer. A bulky residue at the site of cleavage in the peptide substrate reduces the affinity of the enzyme for its substrate. Mn^+2^ primarily improves the catalytic activity by improving the affinity of the enzyme for its substrate, rather than altering the catalytic cleavage rates.

## 2. Materials and Methods

### 2.1. Materials

High purity laboratory grade chemicals were utilized for the experiments. Components for growth media were purchased from Hi-Media (Mumbai, Maharashtra, India), amino acids, dextrose and other chemicals required for study from Sisco Research Laboratories (SRL, Mumbai, Maharashtra, India), Sigma-Aldrich Chemicals (Bangalore, Karnataka, India) and Qualigens (Maharashtra, India), AMC (7-amino-4-methylcoumarin) from Sigma-Aldrich Chemicals (Bangalore, Karnataka, India), glass beads from Unigenetics Instruments Pvt. Ltd, and plasticware from Tarsons (Kolkata, West Bengal, India). All peptide substrates were custom synthesized from GeneScript, USA or Biochain, Biotech Desk Pvt. Ltd. (Hyderabad, India).

### 2.2. Strains and growth conditions

All the strains used in the study were generated from *C. albicans* BWP17 and have been described previously [15] and are listed in Table-I along with their genotypes. The strains were grown in YPD (Yeast Peptone Dextrose) supplemented with uridine. The inoculated primary culture was kept at 30 ℃ at 220 rpm for 16 h. About 2 % of this primary culture was used as an inoculum for 250 ml secondary culture and incubated for 12-14 h under the above-mentioned conditions.

**Table-I:** List of strains used in the study and their genotypes.

| S. No. | Strain | Strain<br>Nomenclature | Genotype | Reference |
| --- | --- | --- | --- | --- |
| 1 | BWP17 | BWP17 | *ura3::imm434/ura3::imm434*<br>*iro1/iro1::imm434*<br>*his1::hisG/his1::hisG arg4/arg4* | [26] |
| 2 | *gpi8Δ* | *GPI8/gpi8* | BWP17- *GPI8/gpi8Δ::HIS1* | [15] |
| 3 | *gpi8Δ/P<sub>GPI8</sub>-*<br>*GPI8* | *GPI8/gpi8-P<sub>GPI8</sub>-*<br>*GPI8* | BWP17-*GPI8/gpi8Δ::HIS1::*<br>*P<sub>GPI8</sub>-GPI8::ARG4* | [15] |

### 2.3. Preparation of post-mitochondrial fraction (PMF)

The preparation of PMF was done as per previously established protocols with minor modifications [16]. Briefly, cells were harvested from 250 ml of *C. albicans* secondary cultures and lysed using acid-washed beads (vortex 22 x 30 s with 30 s on ice after each cycle) in 50 mM Tris-HCl buffer (pH 7.2). The supernatant lysate was aspirated out and centrifuged at 4 °C, first at 3000x g for 15 min and then at 12000x g to obtain the PMF in the supernatant fraction. Protein concentration was estimated by the Bradford assay. Where required, 1-2 ml of the PMF fraction was incubated overnight with EDTA (0.5 mM) at 4 ℃.

### 2.4. Endopeptidase activity assays and steady state kinetics

Unless otherwise specified, the endopeptidase assays were carried out in 100 mM Tris-HCl (pH 7.2). The peptide substrates used were C-terminally linked with 7-amino-4-methyl coumarin (AMC). PMF corresponding to total protein of 0.2 µg was used in a 100 µl reaction volume with varying concentrations of substrate (20-120 µM). To chelate any endogenously bound divalent cation, 0.5 mM EDTA was used. For the metal-dependent kinetics, 5 mM of the required divalent cation was added to the reaction mixture. For pH-dependent assays, the PMF was prepared in either 100 mM sodium acetate buffer (pH 4.5, 5.5, 6.5) or 100 mM Tris-HCl (pH 6.5, 7.2, 7.5). The reactions were normally incubated at 37 ℃ for 6 hours, then centrifuged at 7500xg for 15 min to achieve a clear solution. The supernatant was taken, diluted as required using autoclaved water, and used for spectrofluorimetry.

AMC is a solvent sensitive fluorophore that frequently acts as a reporter in protease assays and has been previously used to study Gpi8 homolog from *T. brucei* [13,17]. When covalently bound to the peptide substrate and excited at 360 nm, AMC has an emission maximum at 400 nm and low quantum yields whereas, but when cleaved from the peptide and released into an aqueous medium, AMC has an emission maximum at 440 nm and significantly higher quantum yields. Hence, to measure product formation, fluorescence emission from free AMC was measured in the range of 380–550 nm after excitation at 360 nm on a Shimadzu RF-5301PC. Slit widths of 3 nm were used for both excitation and emission. Heat-killed controls were taken as controls and their spectra subtracted to obtain the final spectra for the experimental samples. To quantify the endopeptidase activity, fluorescence emission intensity at the peak maximum (440 nm) in the PMF samples was used. Steady state kinetics data were analyzed using Michaelis-Menten plots of 𝑉 versus [𝑆] based on the equation given below:

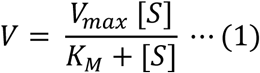

where, 𝑉 is the velocity of the reaction measured by the rate of product formation, 𝑉_𝑚𝑎𝑥_ is the maximum velocity of the reaction, [𝑆] is the concentration of the peptide substrate used in the assay, and 𝐾_𝑀_ is the Michaelis-Menten constant.

All data shown in the manuscript are from independent experiments done thrice in duplicates.

### 2.5. Molecular Dynamics Simulations and Peptide Docking

#### 2.5.1. Model generation for Gpi8_(36-303)_

Model for Gpi8 was generated by AlphaFold Server 3 [18]. The model generated was comparable with PIG-K in human GPIT cryo-EM structure (PDB ID-7WLD). The putative transmembrane regions were removed by using PyMOL 3.1.4.1 3D visualization system to generate Gpi8_(36-303)_. This was done so as to remove the noise that is generated in molecular simulations as the flexibility of these transmembrane helices contribute to higher fluctuations in the simulation data apart from the core portion of Gpi8. In the human GPIT structure the PIG-K subunit alone has a bound Ca^+2^ [11]. The Gpi8 model was therefore aligned with PIG-K using PyMol and the metal ion was introduced in Gpi8_(36-303)_. After that, two other models were generated; the apo model (where the bound Ca^+2^ was removed) and the Mn^+2^-bound model (where the Ca^+2^ was replaced by Mn^+2^). This was done so as to compare the experimental results and to understand the effect of metal ions on overall structural arrangement of the models obtained through molecular simulations.

#### 2.5.2. Molecular dynamics simulation

Molecular Dynamics Simulations were performed using the GROMACS 2022 package [19]. Before MD simulations, the structures were converted to GROMACS format using the pdb2gmx tool. The CHARMM36 force field was employed to generate the protein topology, including parameters for the Ca^+2^ and Mn^+2^ ions present in the structures. The protein structures were then solvated in a cubic solvation box by using the TIP3P water model. Subsequently the whole systems were neutralized by the addition of Na^+^ and Cl^−^ ions. The systems were then energy minimized by using the Steepest Descent minimization algorithm and Verlet cutoff-scheme with maximum number of minimization steps set to 50,000 and maximum force set to less than 10.0 kJ/mol. Thereafter, the energy minimized systems were equilibrated using NVT and NPT methods for 100 ps each, with a time step of 2 fs. The Modified Berendsen Thermostat which was set to 0.1 ps, was used for temperature coupling [20]. The Parrinello-Rahman barostat was used in Pressure coupling and was set to 2 ps. A particle mesh Ewald (PME) [21] algorithm was used to measure long range electrostatic interactions, while the short-range van der Waals and Coulomb interactions were treated using a cut-off scheme, both set to 1.0 nm. The LINCS algorithm [22] was used to constrain all bond lengths. Finally, 100 ns MD simulation was carried out for each structure. The trajectories were analysed to evaluate structural stability and conformational changes, and data analysis was performed using GROMACS. Structural visualization was done using VMD (Visual Molecular Dynamics, stable release 1.9.x) and PyMOL. Trajectory movie for each model was later generated through VideoMach.

Root mean square deviation (RMSD), root mean square fluctuation (RMSF), radius of gyration (Rg), distance and energy plots were calculated using rms, rmsf, gyrate, distance and energy module in GROMACS respectively [20] and plots generated by XMgrace (Grace) 5.1.25.

#### 2.5.3. Peptide docking

The 100 ns structure of all the three models were retrieved in PDB format for protein-peptide docking. HPEPDOCK 2.0 server was used to do flexible peptide docking [23]. The peptide sequence, TIDTNENGS, was used as the substrate and Cys202 was specified as the catalytic site residue. The models generated were further visualized in PyMol and the interacting partners identified.

Hydrogen bond interaction heatmaps between protein and peptide residues were generated using MDAnalysis library in Python [24,25]. The trajectory and topology files were analyzed using the Hydrogen Bond Analysis module with a donor–acceptor distance cut-off < 3.5 Å and an angle cut-off of >150°. Frame-wise hydrogen bond interactions were calculated and converted into percentage occupancy using NumPy. The residue interaction matrix was visualized as a heatmap using Matplotlib.

## 3. Results and Discussion

As mentioned in the Introduction, a cell free assay was established by us for the endopeptidase activity of Gpi8 [15]. This assay used microsomes obtained from wild type (BWP17) [26], as well as heterozygous (*gpi8Δ*) and reintegrant (*gpi8Δ*/P*_GPI8_-GPI8*) strains of *C. albicans*, and a fluorescently labelled seven residue peptide substrate, TIDTNEN-7-amino-4-methyl coumarin (TN7-AMC). Using an improved protocol for the cell free assay that depended on post-mitochondrial fractions (PMF) lacking any divalent cations in the preparation, we obtained steady state kinetic parameters for Gpi8 with different peptide substrates, including TN7-AMC. These results are described below.

### 3.1. Gpi8-specific endopeptidase activity is observed in PMF samples

Cell sub-fractionation and preparation of post-mitochondrial fraction (PMF) was done in the absence of EDTA in Tris-HCl at pH 7.2 as described in the Methods. As can be seen from Figure 1A, the PMF samples from BWP17 cells were active against TN7-AMC. The activity was reduced in *gpi8* and restored in *gpi8Δ*/P*_GPI8_-GPI8*, demonstrating that the activity was specific for Gpi8. The addition of EDTA significantly reduced endopeptidase activity for PMF from all three strains, suggesting that an endogenously bound divalent cation is present in the enzyme (Figure 1B). Addition of Mn^+2^ to the EDTA-treated sample stimulated activity to a larger extent than did the addition of Ca^+2^ (Figure 1C, Figure 1D). The quantification of these data is provided in Figure 1E. Since the PMF samples gave comparable trends in the results as the microsomal assays reported previously [15], the rest of the experiments described here were carried out using freshly prepared PMF.

**Figure 1.**
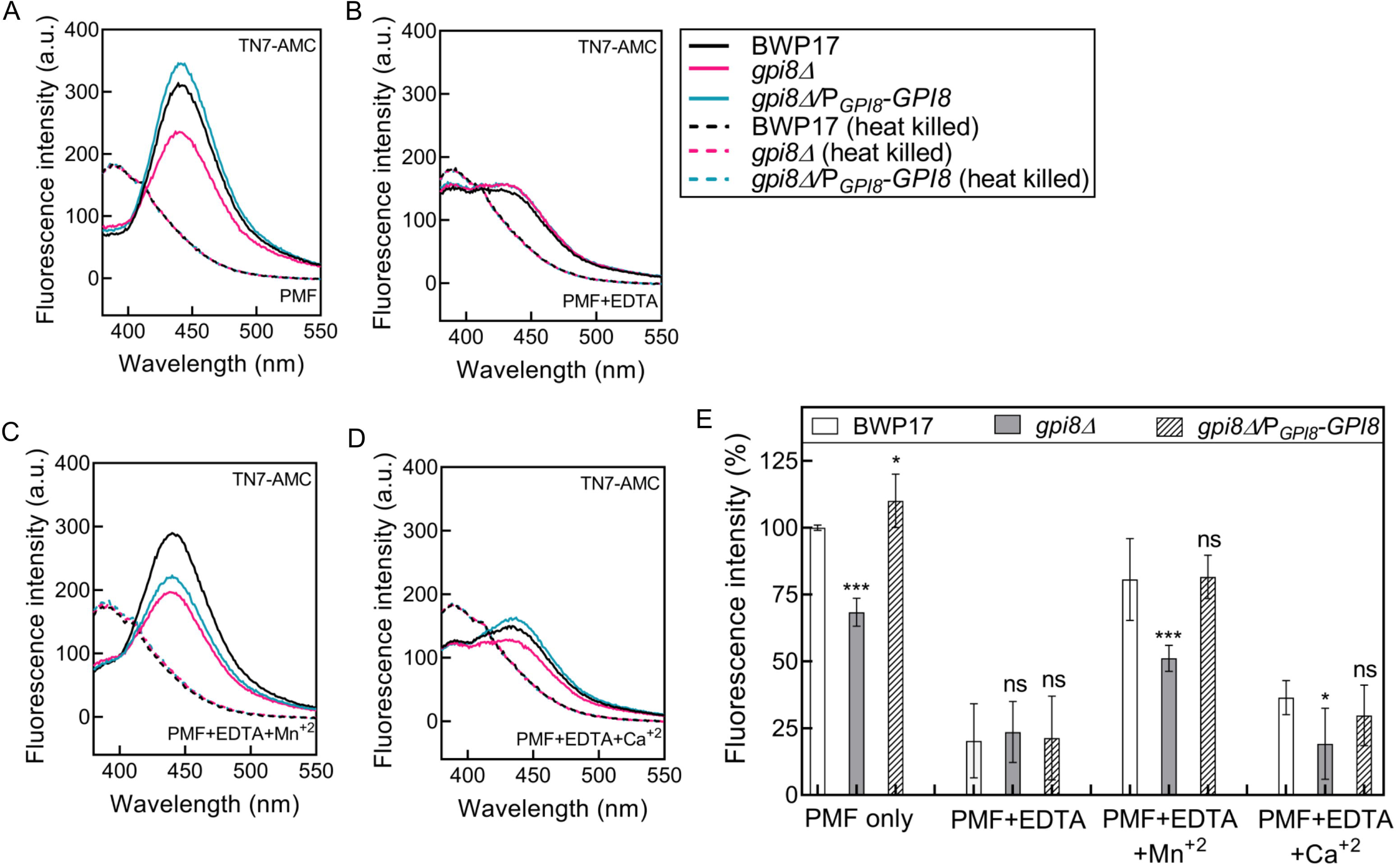
The endopeptidase activity in PMF is Gpi8 dependent. Endopeptidase activity in PMF samples of BWP17, *gpi8Δ* and *gpi8Δ/*P*_GPI8_-GPI8* strains was assayed using TN7-AMC as substrate. The peptide (20 mM) was incubated with PMF samples (each corresponding to 0.2 µg total protein) for 3 h at 37 ℃ in 100 mM Tris-HCl pH 7.2. Free AMC formed in the reaction was detected by excitation at 360 nm and monitoring the fluorescence emission spectra between 380-550 nm using excitation and emission slit widths of 3 nm each. The corresponding heat-killed PMF samples were used as negative controls and their spectra subtracted from that of the experimental samples. Representative spectra shown here correspond to endopeptidase activity in **(A)** freshly prepared PMF, **(B)** EDTA-treated PMF, **(C)** EDTA-treated PMF supplemented with 5 mM MnCl_2_, and **(D)** EDTA-treated PMF supplemented with CaCl_2_. Also shown are the spectra for each of the heat-killed samples. (**E**) The fluorescence emission intensity at the peak maximum (440 nm) were used to quantify the endopeptidase activity in the mutant strains relative to the ‘PMF only’ sample from wild type (BWP17) strain (reported as percentage). The data are averages with standard deviations of three experiments, each done in duplicates, with independent cultures. Black stars indicate the significance of data relative to BWP17. Note: a.u. implies arbitrary units. The symbols *, and ***, correspond to P-values ≤ 0.05, and ≤ 0.0005, respectively, while ‘ns’ implies non-significant, relative to BWP17.

### 3.2. The endopeptidase activity is optimum at pH 7.2

In order to test the optimum pH for Gpi8 endopeptidase activity, the activity assays were carried out in buffers of different pH. As can be seen from (Figure 2A), the activity of freshly prepared PMF was a maximum at pH 7.2. Treatment with EDTA resulted in drastically reduced activity (Figure 2B) but when supplemented with Mn^+2^ the activity once again peaked at pH 7.2 (Figure 2C). Clearly, the divalent cation did not alter the optimum pH of the enzyme, suggesting that the metal had no role in creating a nucleophile for catalysis. Also, given that the enzyme activity is comparable at pH 6.5 in both Tris-HCl and sodium acetate buffers (Figure 2D), it may also be inferred that the choice of buffer did not affect the results.

**Figure 2.**
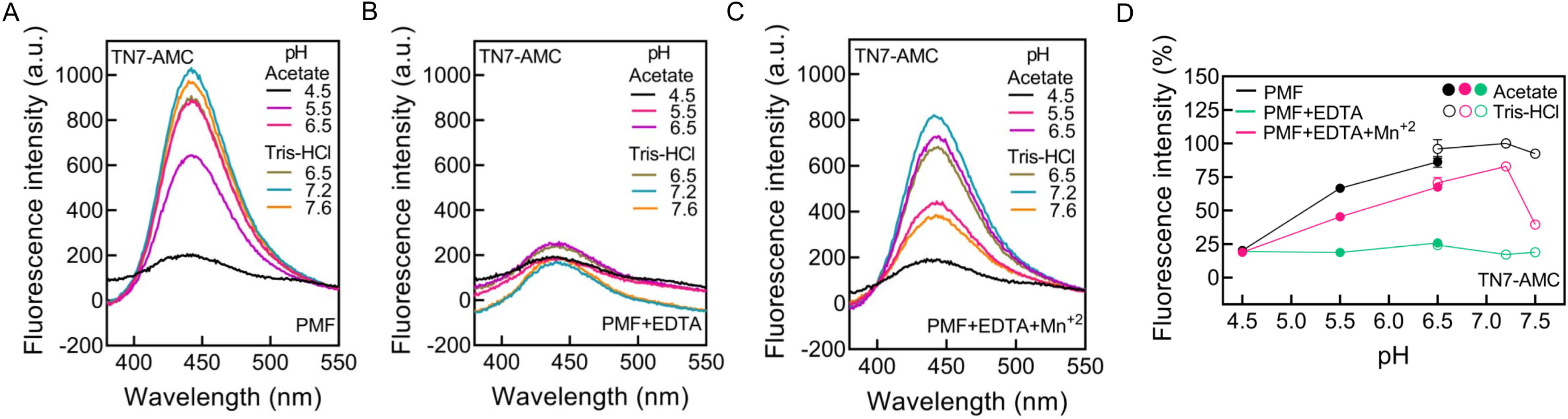
The endopeptidase activity is maximum at pH 7.2. TN7-AMC (20 mM) was incubated for 6 h at 37 ℃ with PMF samples (corresponding to 0.2 µg total protein) prepared from BWP17 in different pH conditions (100 mM acetate buffer at pH 4.5, 5.5 and 6.5) and (100 mM Tris-HCl buffer at pH 6.5, 7.2 and 7.5). The product (AMC) formed was monitored by excitation at 360 nm and recording fluorescence emission spectra between 380-550 nm with excitation and emission slit widths of 3 nm each. Representative spectra at different pH are shown for **(A)** PMF only, **(B)** PMF treated with EDTA, **(C)** PMF treated with EDTA and then supplemented with 5 mM MnCl_2_. The spectra of the corresponding heat-killed PMF samples, used as negative controls, were subtracted from that of the experimental samples. **(D)** Scatter plots obtained using the fluorescence emission intensity at 440 nm to quantify the endopeptidase activity in freshly prepared PMF, freshly prepared PMF treated with EDTA, and freshly prepared PMF treated with EDTA followed by addition of 5 mM MnCl_2_ in the different pH conditions relative to that for PMF alone at pH 7.2 (reported as percentage). The data are averages with standard deviations of three experiments, each done in duplicates, with independent cultures. Note: a.u. implies arbitrary units.

### 3.3. The endopeptidase activity is stimulated by specific divalent cations

An expanded panel of divalent cations was then tested at pH 7.2 using TN7-AMC as substrate to confirm the divalent cation specificity of Gpi8 in the PMF sample from BWP17 strain. While Mg^+2^ and Zn^+2^ showed no significant stimulation of endopeptidase activity both Cu^+2^ and Ni^+2^ showed a small inhibition of activity; Ca^+2^ enabled a small stimulation while Mn^+2^ showed a significantly higher stimulation of endopeptidase activity (Figure 3). As additional controls, the fluorescence of the peptide substrate alone (Figure S1A) as well as the product (AMC) alone (Figure S2) were monitored in the absence and presence of EDTA, as well as in the presence of either Mn^+2^ or Ca^+2^. No significant difference was seen in the spectra at 440 nm in any of these conditions, suggesting that the changes in fluorescence intensities seen in Figure 3 were not related to either EDTA or the divalent cations binding directly to either of them.

**Figure 3.**
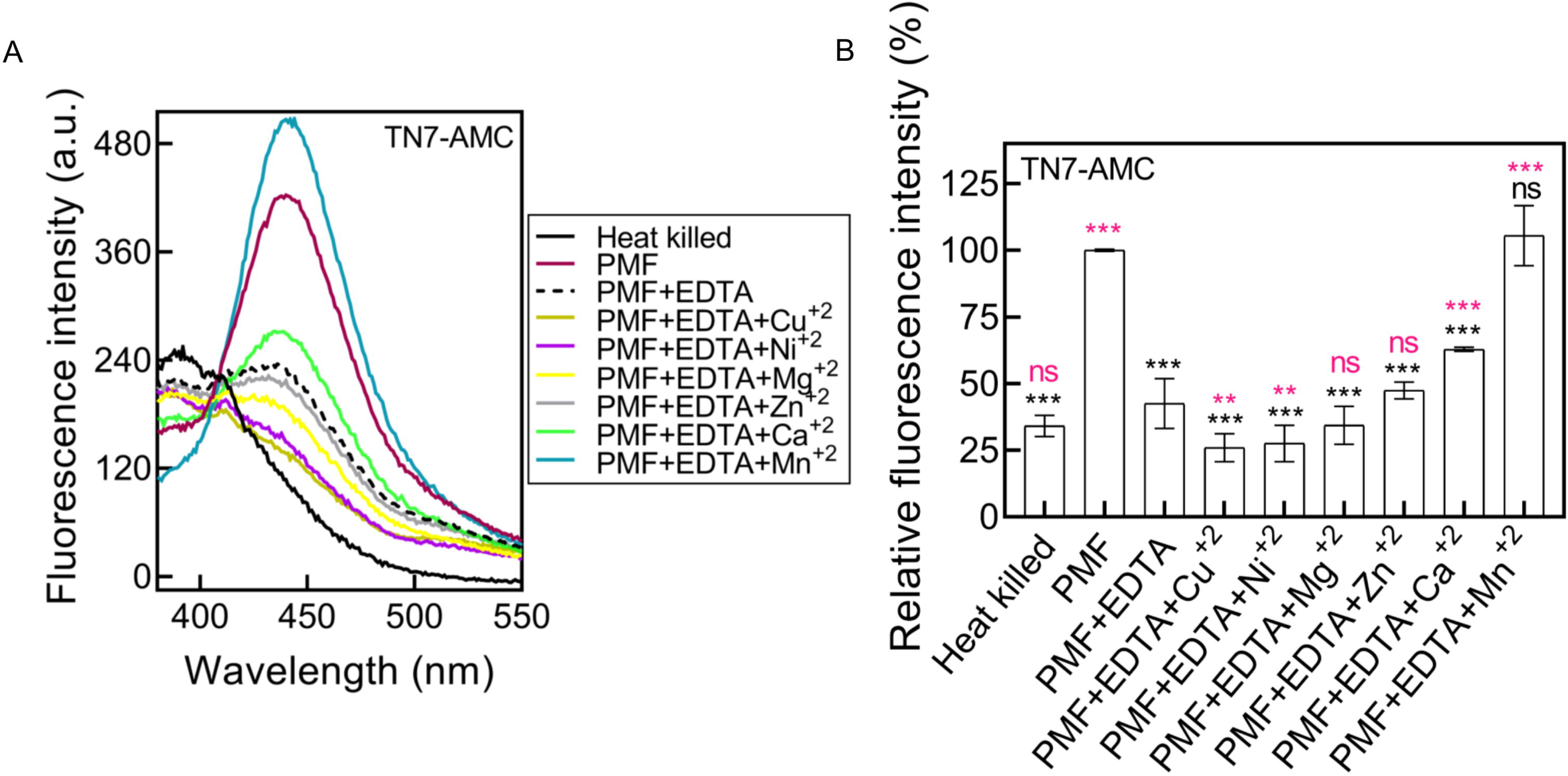
Divalent cations stimulate Gpi8 activity. Freshly prepared PMF samples from BWP17 strain were incubated with EDTA and then supplemented with 5 mM solutions of different divalent cations as shown in the figure. PMF, PMF treated with EDTA, and PMF-treated with EDTA supplemented with different divalent cations were incubated with TN7-AMC (20 mM) for 6 h at 37 ℃ in 100 mM Tris-HCl (pH 7.2). The product (AMC) formed was detected by excitation at 360 nm and emission recorded between 380-550 nm with slit widths of 3 nm for excitation and emission. **(A)** Spectra showing AMC formed in the absence and presence of EDTA as well as after supplementation with the divalent cations. The spectra obtained from peptide incubated with heat-killed PMF sample is also shown. **(B)** Bar graph showing relative endopeptidase activity in the same samples relative to that in the freshly prepared PMF sample (reported as percentage). The data are averages with standard deviations of three experiments, each done in duplicates, with independent cultures. Black stars represent significance of the data relative to PMF alone, while magenta stars represent significance of the data relative to the EDTA-treated PMF. Note: a.u.; arbitrary units. The symbols *, ** and ***, correspond to P-values ≤ 0.05, ≤0.005, and ≤ 0.0005, respectively, while ‘ns’ implies non-significant.

### 3.4. Steady state kinetics of the endopeptidase activity in a cell free system

To determine the steady state kinetic parameters, it was necessary to determine the concentration of the product formed. So, every time an experiment was performed, we also obtained in parallel a standard curve for AMC using known concentrations of it and used it to estimate the concentration of the product formed in the assay (Figure S3).

Peptide substrates for the assays were custom synthesized. The peptides, VN9 (VPTIDTNEN) and TN15 (TDSSSSVPTIDTNEN), are both N-terminal elaborations of TN7 (TIDTNEN) based on the signal sequence residues of Hyr1, a GPI-anchored protein of *C. albicans* while TN4 (TNEN) is a N-terminally truncated version of TN7 (Table-II). The terminal amino acid in all the four peptides is an Asn (Asn 895 in Hyr1), based on the predicted cleavage site or the ω site, at which GPI attachment should occur in the cell as per three different predictive softwares: NetGPI-1.1 [27–29]. TP7 (TIDTNEP), on the other hand, is a site-specific mutant of TN7 in which Asn at the ω site is replaced by Pro (Table-II). The fluorophore, AMC, was C-terminally linked to the peptide in all cases. Steady state kinetic assays were performed with each of peptide substrates and K_m_ as well as V_max_ values determined.

**Table-II.**
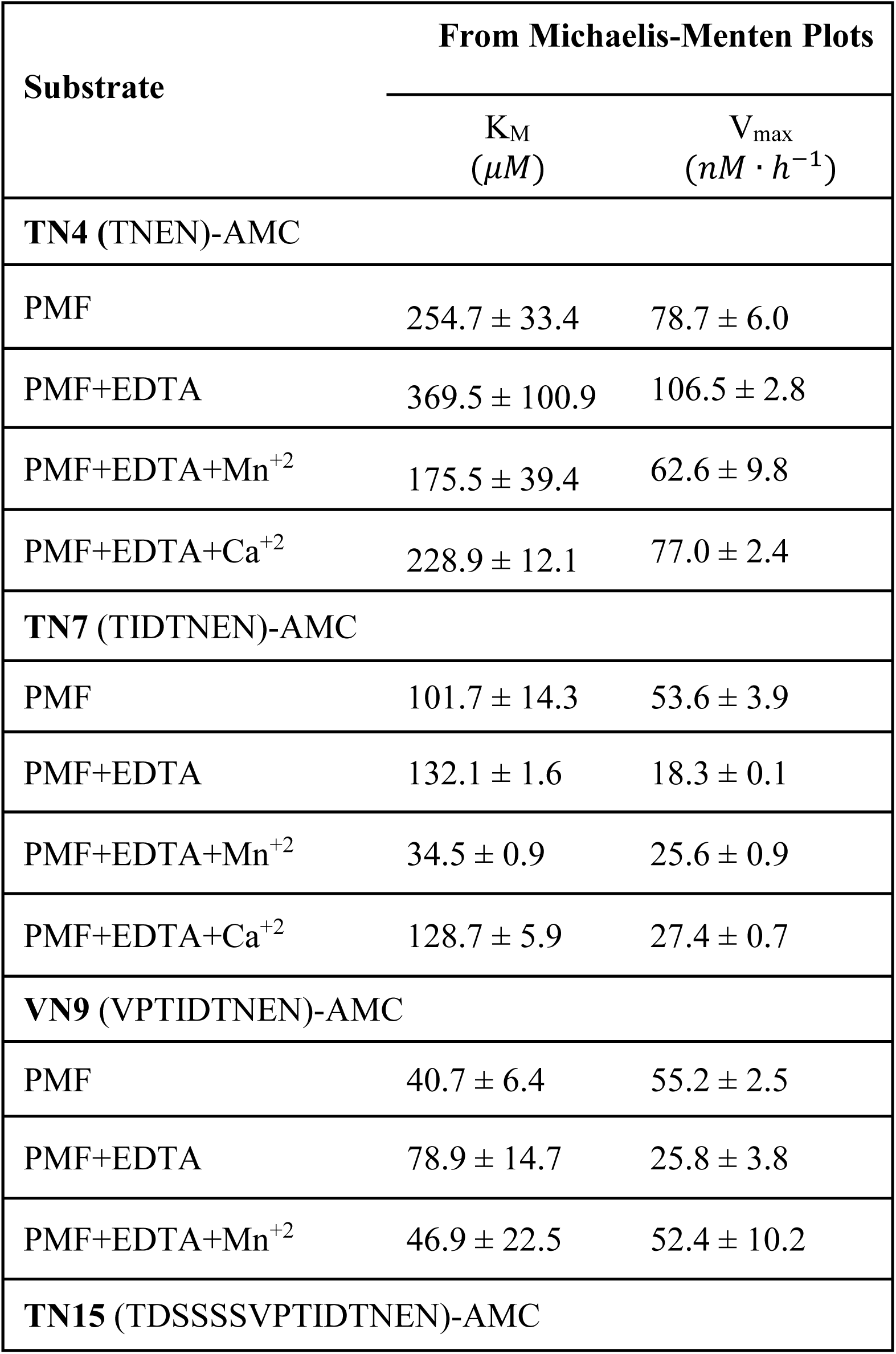
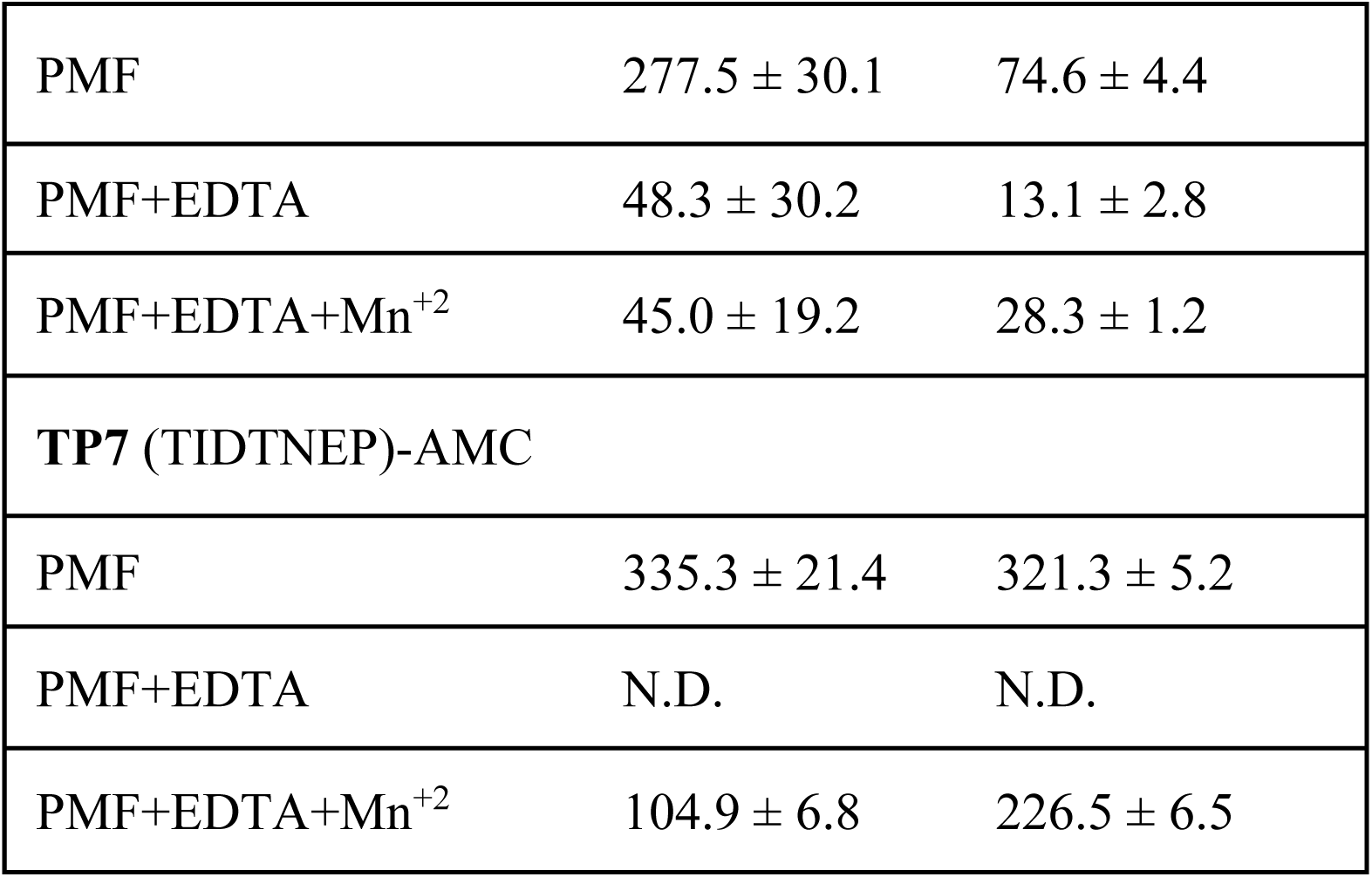
Steady state kinetics parameters of CaGpi8 with respect to different peptide substrates under different experimental conditions. PMF corresponding to total protein of 0.2 µg was used in a 100 µl reaction volume with varying concentrations of substrate (20-120 µM). Tris-HCl buffer (100 mM; pH 7.2) was used for the assay. Where mentioned, EDTA was used at a concentration of 0.5 mM. MnCl_2_ and CaCl_2_ were used at concentrations of 5 mM. The values of K_M_ and V_max_ were obtained from Michaelis-Menten plots of the data.

It may be noted that in all cases, the growth conditions of the strain and total amount of protein used in the assays were kept constant. The PMF was always freshly prepared for each experiment. Data from assays using heat-killed PMF samples for each concentration of peptide were subtracted to correct for any non-specific contribution from background. Also, as with TN7-AMC (Figure S1A), the fluorescence spectrum of none of the other peptide substrates alone in the absence or presence of EDTA, was significantly affected by the presence of either Mn^+2^ or Ca^+2^ (Figure S1B-E). Further, in order to minimise errors, a control assay with TN7-AMC was carried out in parallel with the assays for any of the other peptides.

#### 3.4.1. Role of the divalent cation

Steady state kinetic assays were first performed with TN7-AMC as substrate (Figure 4A (i)-(iv)), the substrate we used in our previous studies, and the Michaelis-Menten curves were plotted (Figure 4A(v)). The representative spectra for freshly prepared PMF (Figure 4A(i)) show that the endopeptidase activity increases in a substrate concentration-dependent manner. Addition of EDTA greatly reduces the endopeptidase activity (Figure 4A(ii)) while the addition of Mn^+2^ significantly stimulates it (Figure 4A(iii)).

**Figure 4.**
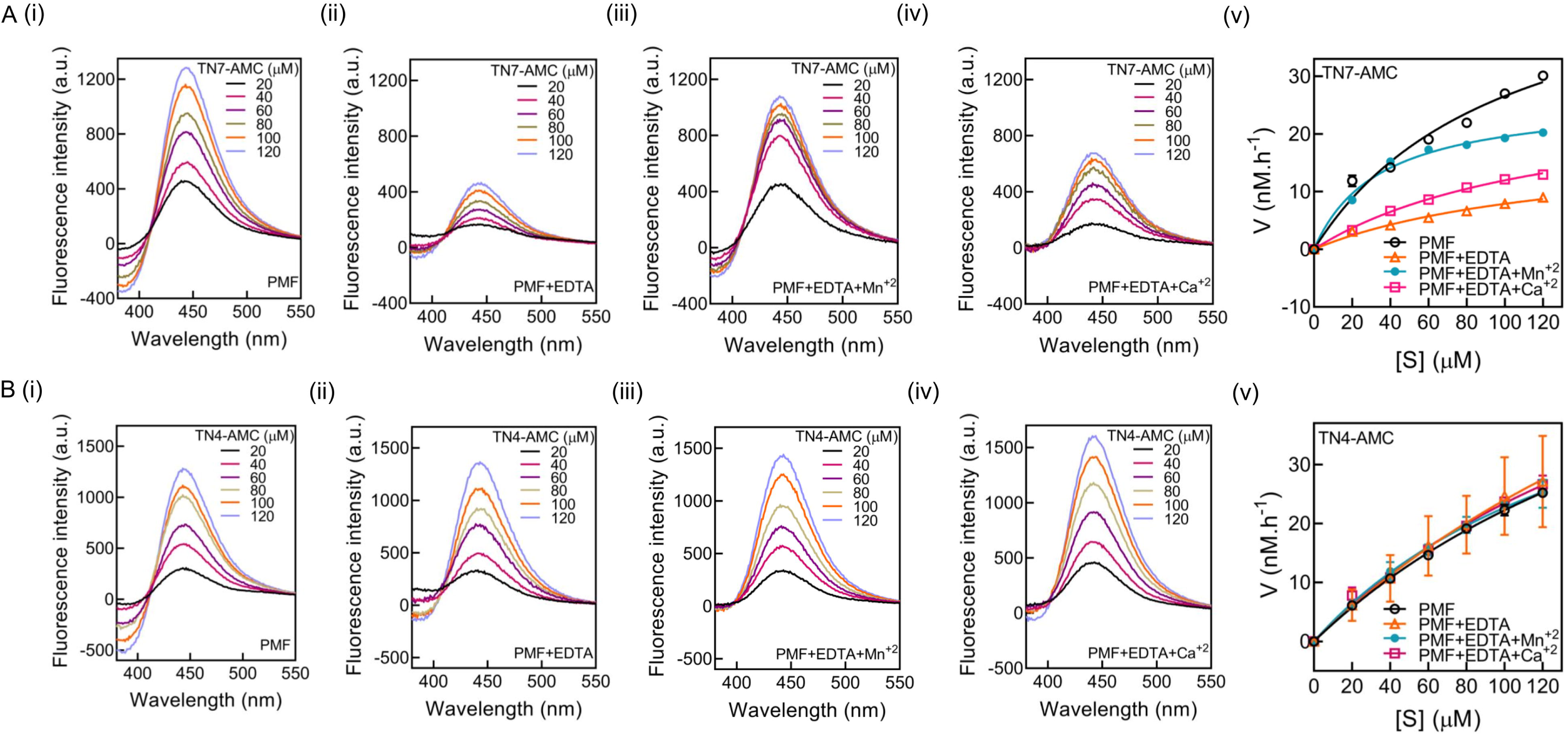
Steady state kinetics of the endopeptidase activity of Gpi8 using (A) TN7-AMC and (B) TN4-AMC. Fluorescence emission spectra of free AMC produced after different concentrations of TN7-AMC (20 µM to 120 µM) were incubated for 6 h, at 37 ℃ in 100 mM Tris-HCl (pH 7.2) in the presence of **(i)** PMF, **(ii)** EDTA-treated PMF, **(iii)** EDTA-treated PMF stimulated by 5 mM MnCl_2_ and **(iv)** EDTA-treated PMF supplemented with 5 mM CaCl_2_. The samples were excited at 360 nm with 3 nm slit widths for excitation and emission. Fluorescence intensity at 440 nm was used to estimate the concentration of AMC formed, based on a standard curve for AMC alone (representative plot shown in Figure S3). This was then used to plot Michaelis-Menten curves shown in **(v)** from which K_M_ and V_max_ values were obtained (Table-II). The data are averages with standard deviations of three experiments, each done in duplicates, with independent cultures. Note: a.u.; arbitrary units.

The steady state kinetics data for this substrate provided in Table-II are very informative. Freshly prepared PMF, containing no EDTA, had a K_M_ of 101.7 ± 14.3 µM for this substrate while addition of EDTA increases this to 132.1 ± 1.6 µM. Addition of 5 mM Mn^+2^ to the EDTA-treated sample, brings down the K_M_ to 34.5 ± 0.9 µM. A few points are immediately obvious. The addition of EDTA appears to extract an endogenously bound divalent cation, and reduce the affinity of the enzyme for its substrate (increases K_M_). Addition of Mn^+2^ to this EDTA-treated sample improves the affinity of the enzyme for its peptide substrate (reduces K_M_), indicating that the divalent cation plays a role in substrate binding. Substrate affinity is also improved upon addition of Mn^+2^ as compared to that obtained with freshly prepared PMF alone, suggesting either that the endogenously bound cation is different, or that the process of preparing the PMF causes at least some of the GPIT complexes present in the sample to lose their bound divalent cations. The V_max_ for TN7-AMC also reduces approximately three-fold from 53.6 ± 3.9 nM.h^−1^ to 18.3 ± 0.1 nM.h^−1^ upon addition of EDTA, indicating that the loss of an endogenously bound divalent cation affects the catalytic activity as well. Expectedly, then, addition of Mn^+2^ improves V_max_ to 25.6 ± 0.9 nM.h^−1^.

As can be seen from Figure 4A (iv)-(v) and Table-II, the addition of Ca^+2^ does not stimulate the endopeptidase activity to the same extent as Mn^+2^ nor does it improve the K_M_ value to the same extent. In fact, in the presence of Ca^+2^, the K_M_ (128.7 ± 5.9 µM) is comparable to that of the EDTA-treated sample but the V_max_ (27.4 ± 0.7 nM.h^−1^) is comparable to that obtained in the presence of Mn^+2^. Given that Mn^+2^ has a significantly smaller ionic radius as compared to Ca^+2^, it is possible to speculate that the smaller divalent cation induces a greater conformational change in the enzyme that enables better substrate binding. In other words, the choice of the divalent cation appears to dictate substrate binding but not catalytic rates.

Similar trends are obtained when we use a shorter peptide, TN4-AMC, in the assay. It is worth noting that the abstraction of the endogenously bound divalent cation does not cause a drastic loss in activity in the case of this shorter peptide (compare Figure 4B(ii) with Figure 4B(i)). Thus, for a shorter peptide, the metal-induced stabilization does not appear to contribute much to the binding. As a result, stimulation of endopeptidase activity by Mn^+2^ and Ca^+2^ are also lesser (Figure 4B(iii)-(iv)).

This is reflected in the Michaelis-Menten plots too (Figure 4B(v)). The steady state kinetic parameters obtained from these plots (Table-II) suggest that treatment with EDTA leaves the K_M_ relatively unchanged within the limits of error (from 254.7 ± 33.4 µM to 369.5 ± 100.9 µM). The addition of Mn^+2^ only marginally improves substrate affinity (K_M_ =175.5 ± 39.4 µM) as does the addition of Ca^+2^ (K_M_ = 228.9 ± 12.1 µM).

The V_max_ of the enzyme in freshly prepared PMF is 78.7 ± 6.0 nM.h^−1^ for TN4-AMC, while in the presence of EDTA it is 106.5 ± 2.8 nM.h^−1^. This improvement in catalytic rates is probably due to the ease of leaving of the product from the active site. Perhaps abstraction of the divalent cation causes an opening up of the catalytic pocket. Addition of Mn^+2^ brings the V_max_ down to 62.6 ± 9.8 nM.h^−1^, and the addition of Ca^+2^ to 77.0 ± 2.4 nM.h^−1^.

#### 3.4.2. Importance of substrate length

As seen above, under similar assay conditions, the enzyme in the PMF has ∼2.5-fold weaker affinity for TN4-AMC as compared to TN7-AMC (Table-II). A smaller peptide like TN4-AMC probably makes fewer contacts with the substrate binding site, which probably explains the higher K_M_. This led us to examine whether there was an optimum substrate length required for catalysis by Gpi8.

For this purpose, we used two longer peptide substrates, VN9-AMC and TN15-AMC, in the assay (Figure 5 A&B; Table-II). The enzyme has ∼2.5-fold higher affinity for VN9-AMC (K_M_ = 40.7 ± 6.4 µM) relative to TN7-AMC. Upon addition of EDTA, the affinity for VN9-AMC reduces by roughly 2-fold as does the catalytic rates (K_M_ = 78.9 ± 14.7 µM; V_max_ = 25.8 ± 3.8 nM.h^−1^), while addition of Mn^+2^ to the EDTA-treated sample restored substrate affinity and catalytic rates (K_M_ = 46.9 ± 22.5 µM; V_max_ = 52.4 ± 10.2 nM.h^−1^).

**Figure 5.**
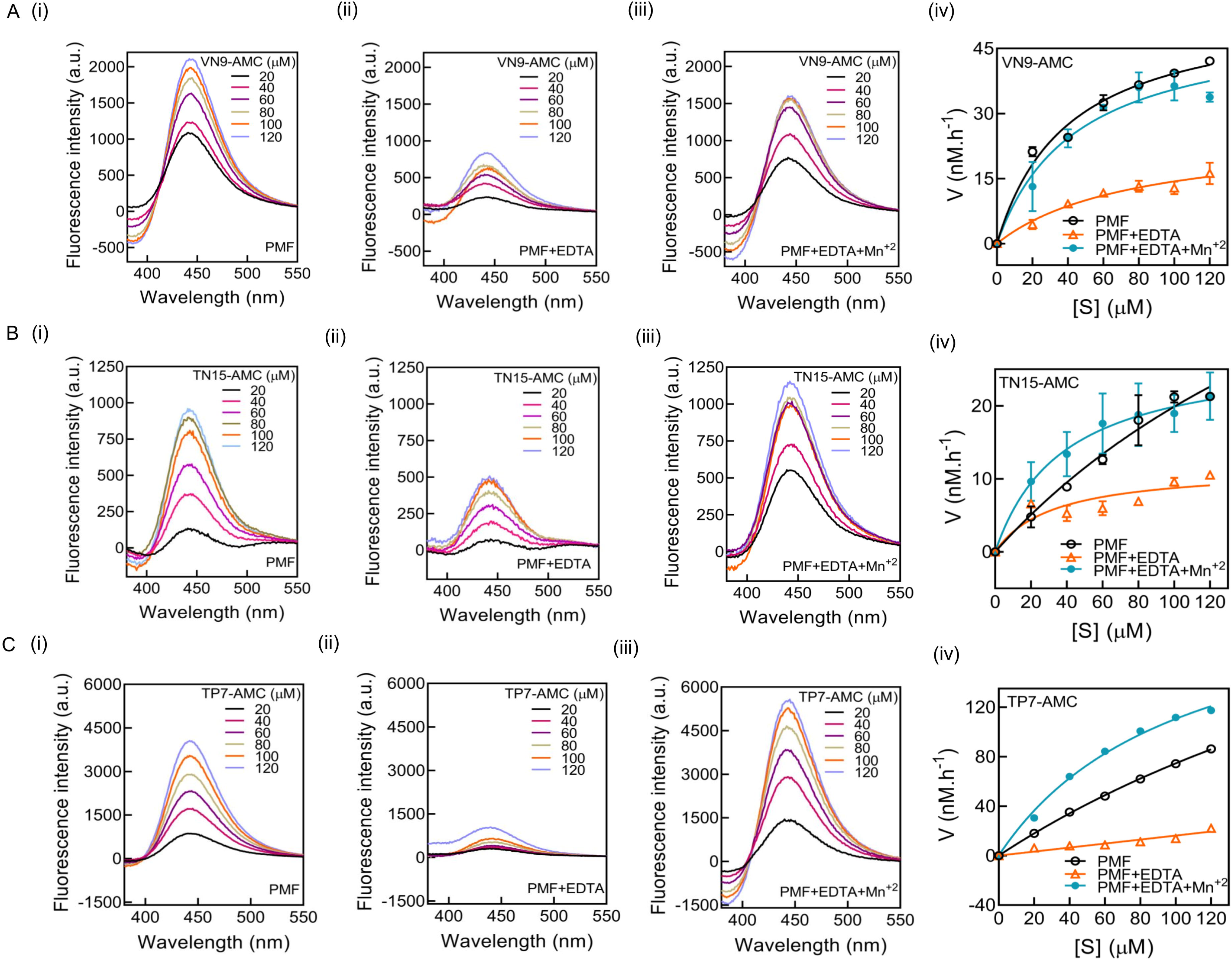
Steady state kinetics of the endopeptidase activity using (A) VN9-AMC, (B) TN15-AMC, and (C) TP7-AMC. Fluorescence emission spectra of free AMC produced after different concentrations of the peptide substrates (20 µM to 120 µM) were incubated for 6 h, at 37 ℃ in 100 mM Tris-HCl (pH 7.2) in the presence of **(i)** PMF, **(ii)** EDTA-treated PMF, and **(iii)** EDTA-treated PMF supplemented with 5 mM MnCl_2_. The samples were excited at 360 nm with 3 nm slit widths for excitation and emission. Fluorescence intensity at 440 nm was used to estimate the concentration of AMC formed, based on a standard curve for AMC alone (representative plot shown in Figure S3). This was then used to plot Michaelis-Menten curves shown in **(iv)** from which K_M_ and V_max_ values were obtained (Table-II). The data are averages with standard deviations of three experiments, each done in duplicates, with independent cultures. Note: a.u. represents arbitrary units.

In the case of TN15-AMC, the enzyme has ∼2.5-fold lower affinity (K_M_ = 277.5 ± 30.1 µM) relative to TN7-AMC (K_M_ =101.7 ± 14.3 µM). The addition of EDTA reduces the catalytic activity to such a large extent, that it was not possible to reliably estimate K_M_, and the errors are large. However, addition of Mn^+2^ to the EDTA-treated sample improved both substrate affinity and catalytic rates (K_M_ = 45.0 ± 19.2 µM; V_max_ = 28.3 ± 1.2 nM.h^−1^).

Thus, it may be possible to infer the following: 1) the length of the substrate is an important factor determining substrate binding affinity and, 2) that 7-mer to 9-mer peptides probably fit optimally within the catalytic pocket for efficient cleavage by the enzyme.

#### 3.4.3. Relevance of the size of the ω site residue

To study whether our assay would be sensitive to alterations in the size of the residue at ω site we used TP7-AMC, a 7-mer peptide substrate where AMC is attached to Pro rather than Asn as in TN7-AMC (Figure 5C; Table-II). Freshly prepared PMF exhibited nearly 3-fold weaker affinity for TP7-AMC (K_M_ = 335.3 ± 21.4 µM) as compared to TN7-AMC (101.7 ± 14.3 µM). The more bulky Pro appears to fit poorly into the active site and reduces affinity. However, V_max_ for TP7-AMC (321.3 ± 5.2 nM.h^−1^) is better as compared to TN7-AMC (53.6 ± 3.9 nM.h^−1^). It is possible that the poorer fit makes it easier for the product to leave the catalytic pocket, thus improving catalytic rates. In this case too, the addition of EDTA results in very low activity, as a result of which the steady state parameters could not be reliably obtained. The addition of Mn^+2^ stimulates activity and improves K_M_ to 104.9 ± 6.8 µM, but the V_max_ remains ∼1.5-fold lower than for the enzyme in freshly prepared PMF (226.5 ± 6.5 nM.h^−1^ versus 321.3 ± 5.2 nM.h^−1^).

### 3.5. Molecular dynamics simulation studies

#### 3.5.1. The model of Gpi8 compared will with that for human PIG-K

The model for Gpi8 was generated by AlphaFold Server 3 and was found to compare well with PIG-K in human GPIT cryo-EM structure (PDB ID-7WLD) with an RMSD of 0.757 (Figure S4) [11]. The N-terminal transmembrane regions were removed by using PyMOL 3.1.4.1 3D visualization system to generate the soluble form of the protein, Gpi8_(36-303)_, which contains the putative catalytic domain as well.

#### 3.5.2. Divalent cations stabilize the soluble domain of Gpi8

The RMSD data obtained for the truncated protein indicate greater structural changes in the apo form as compared to the Mn^+2^-bound and Ca^+2^-bound forms, suggesting that the divalent cations provide greater rigidity to the overall structure of the soluble domain (Figure 6A). The RMSD data for the Ca^+2^-bound model suggests that this model equilibrated better over the entire run. Further, it appears to be the most rigid, or having the least RMSD, in the overall structure over the entire duration of the simulation. Based on the models generated, Asp75, Ile78 and Asp116 appear to be the metal-binding residues for both Ca^+2^ and Mn^+2^ (Figure 6B (i)-(ii)). Homologous residues are also conserved in other organisms (Asp79, Ile82 and Asp120 in human PIG-K; Asp72, Ile75 and Asp113 in *S. cerevisiae* Gpi8 and Asp63, Ile66 and Asp106 in *T. brucei* PIG-K, respectively) (Figure 6C).

**Figure 6.**
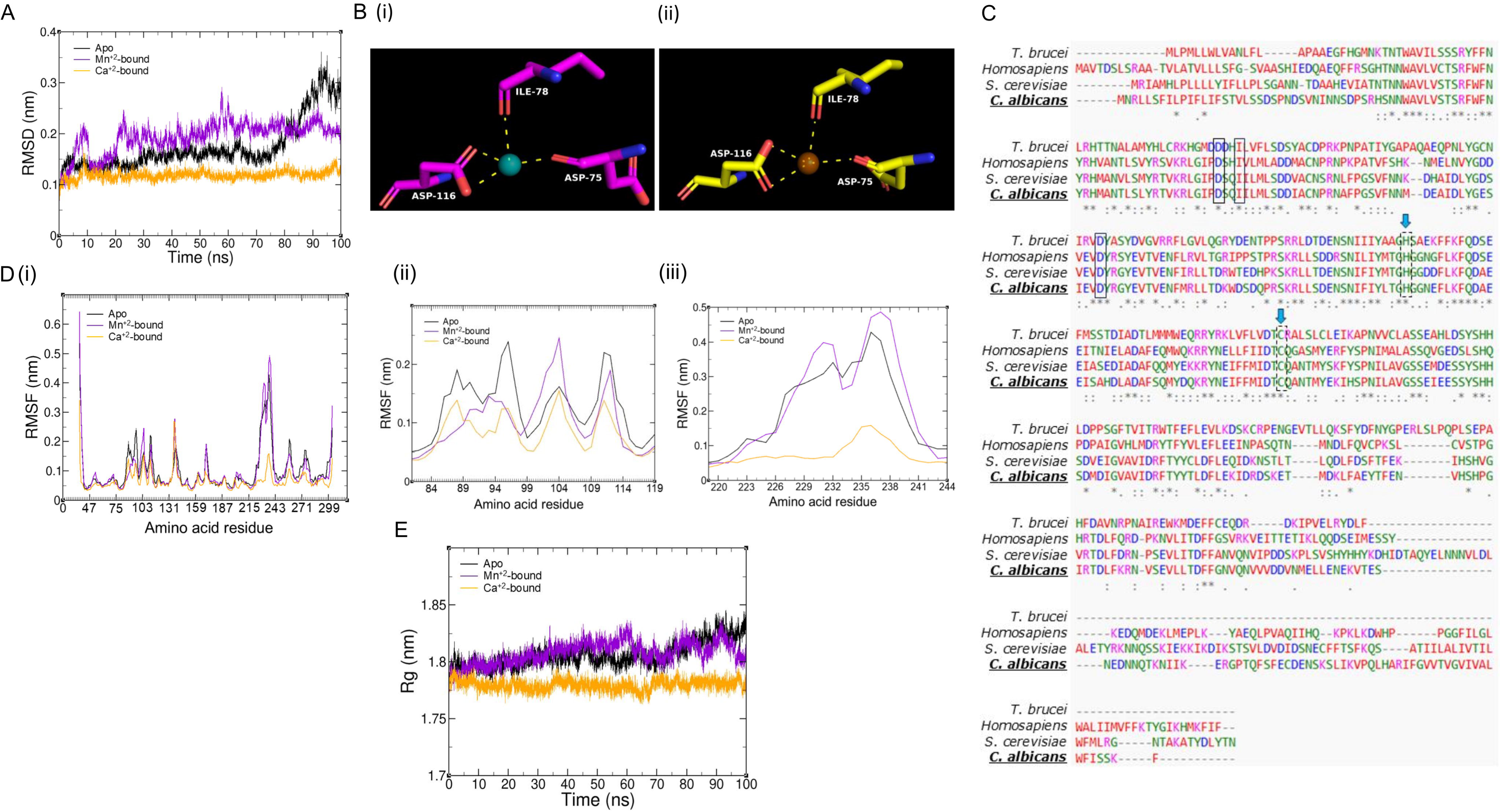
*In silico* analysis suggests that divalent cations stabilize soluble domain of Gpi8. **(A)** Root Mean Square Deviation (RMSD) showing structural deviations between apo and metal bound forms of Gpi8_(36-303)_ after a molecular simulation run of 100 ns. **(B)** Snapshot showing metal binding residues of Mn^+2^– and Ca^+2^–bound Gpi8_(36-303)_ from the 100 ns frames. Mn^+2^ is represented in cyan and Ca^+2^ in orange. Metal coordination residues are represented as magenta-coloured sticks for **(i)** Mn^+2^-bound Gpi8_(36-303)_ and as yellow-coloured sticks for **(ii)** Ca^+2^-bound Gpi8_(36-303)_. **(C)** Multiple sequence alignment of Gpi8 from *C. albicans*, *Homosapiens*, *S. cerevisiae* and *T. brucei* as obtained from Clustal Omega. Metal coordination residues are enclosed in closed rectangular box while catalytic dyads are enclosed in dotted boxes represented by blue arrows. **(D)** Root Mean Square Fluctuation (RMSF) at each residue in Gpi8_(36-303)_. **(i)** RMSF at each residue in apoGpi8_(36-303)_, Mn^+2^-bound Gpi8_(36-303)_, and Ca^+2^-bound Gpi8_(36-303)_ show significant differences between the three models. The residual fluctuations are particularly prominent in the regions 80-119, 130-144, 169-176, 219-244, 244-269 and 269-280. **(ii)** Enlarged view of the RMSF plot of structural fluctuations happening near the metal binding site. **(iii)** Enlarged view of the RMSF plot of structural fluctuations happening near the loop region (219-244) close to catalytic pocket. **(E)** Radius of Gyration (Rg) plots show structural compactness in the metal bond form relative to apoGpi8_(36-303)_. The plots were generated by XMgrace.

The root mean square fluctuations (RMSF) at different residues of Gpi8_(36-303)_ were also plotted (Figure 6D(i)-(iii)). From these plots it is evident that there are large changes in amino acid positions in all 3 models, but the apo form shows greater fluctuations and the Ca^+2^-bound form the least. These fluctuations are particularly prominent at amino acid positions that are involved in metal coordination (Figure 6D(ii)) and at a flexible loop region (219-244) that is proximal to the catalytic pocket (Figure 6D(iii)). It is also worth mentioning that only small fluctuations are observed at the catalytic site residues, Cys202 and His160 *per se*. Further, the metal coordination remains stable and they remain bound to the protein throughout the simulation.

The data for radius of gyration (Rg) shows that all the three models are well folded and the apo form does not unfold during the entire simulation period (Figure 6E). Thus, it appears that a divalent cation is not crucial for protein stability. However, the metal does enable compaction of the protein structure.

#### 3.5.3. Energy calculations show that metal binding enhances stability of Gpi8_(36-303)_

For further understanding the implications of these changes in structure between the three forms, the potential energy (the sum of all the bonded and non-bonded interactions, which reflects the overall stability of the system) and Lennard Jones (LJ) short range (SR) interactions were calculated from the molecular simulations. From the potential energy data (Figure 7A (i)-(ii); Table-III), it can be noted that the metal bound models have lower mean potential and are stabilized as compared to the apo form. Further, the LJ-SR energy values remain comparable in all 3 cases (Figure 7B; Table-III) indicating that proper hydrophobic packing and overall steric fit are not affected by the divalent cations. Therefore, the overall core packing remains largely intact even in the absence of a divalent cation.

**Figure 7.**
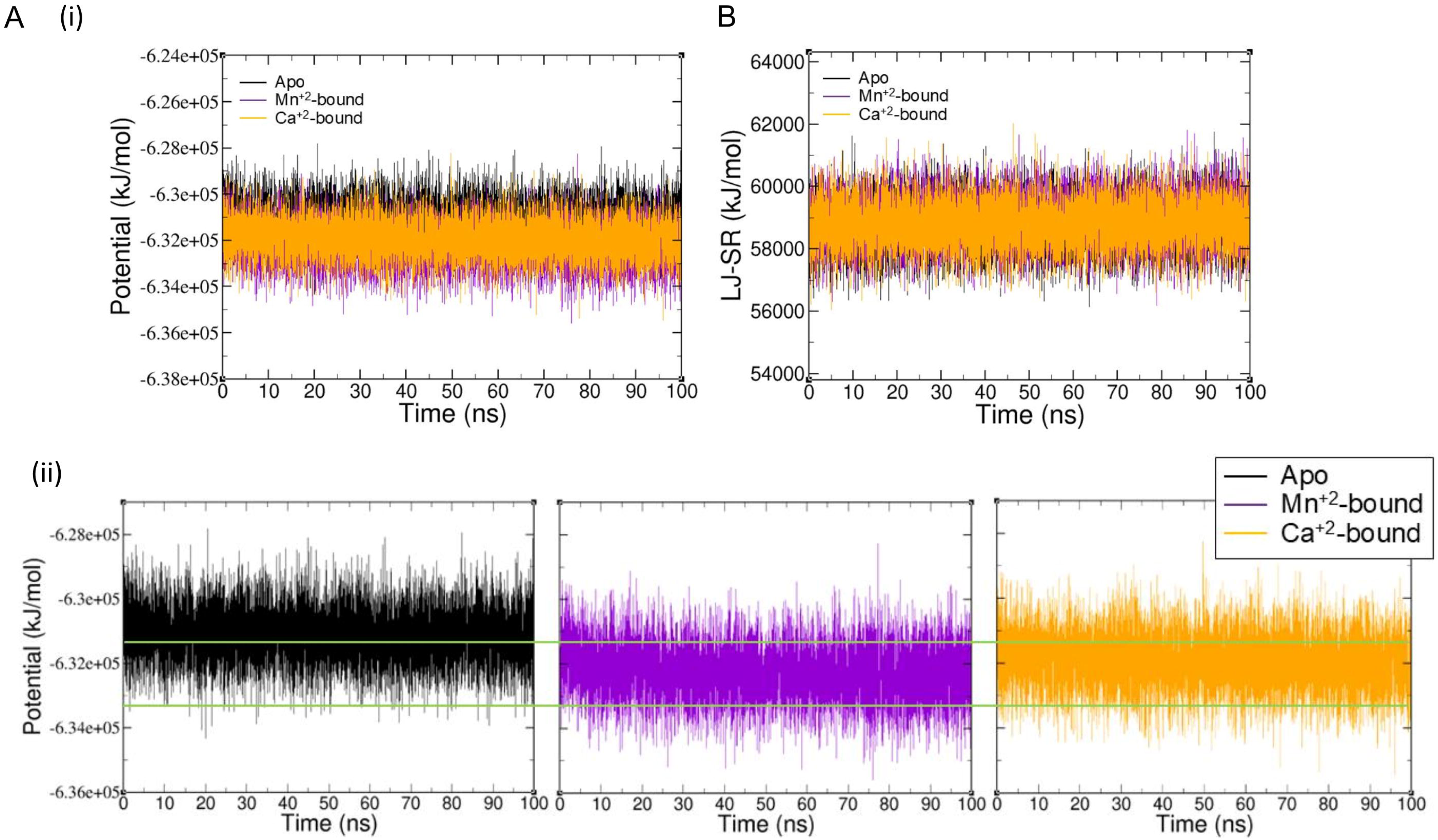
Potential energy plot show differences in energy states of apo and metal bound forms of Gpi8_(36-303)_. **(A)** Long-range electrostatic interactions for **(i)** all the three models during the 100 ns molecular dynamics simulation run and **(ii)** for apoGpi8_(36-303)_, Mn^+2^-bound Gpi8_(36-303)_, and Ca^+2^-bound Gpi8_(36-303)_ individually. The green line is added for easy comparison between the trendlines for each model. **(B)** LJ-SR (Lenard Jones-Short Range) interactions are comparable between apo and metal bound forms of Gpi8_(36-303)_, suggesting similar hydrophobic packing and steric effects.

**Table III.**
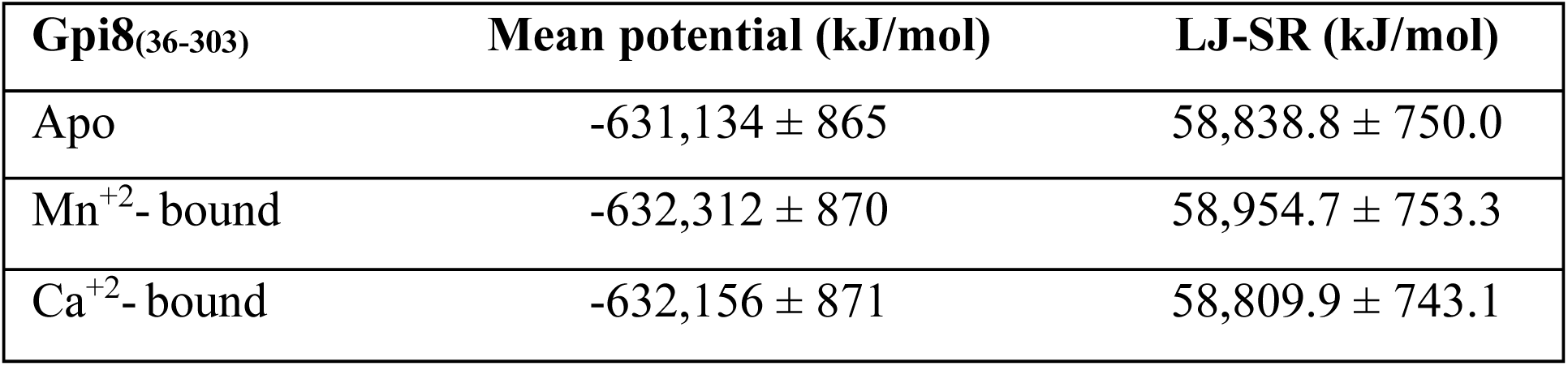
Energy of the apo and metal bound models. The mean potential energy and Lennard Jones short range (LJ-SR) interactions as calculated from the molecular dynamic simulations.

#### 3.5.4. Loop fluctuations and active site dynamics within the models

A trajectory movie, with 1 ns interval across the entire run, was generated for each model from centered MD simulations using VMD to visualize loop and active-site dynamics. For comparison, the 100 ns structures of Gpi8_(36-303)_– apo, Mn^+2^-bound, and Ca^+2^-bound forms – were retrieved, aligned and analyzed through PyMOL and are shown in (Figure 8 A-D). From these structures it is obvious that most of the structural changes observed are around the flexible loops enveloping the protein rather than within the core. This correlates with the Rg and RMSF data discussed above, indicating that the protein structure remains intact but loop rearrangements are observed within the structures.

**Figure 8.**
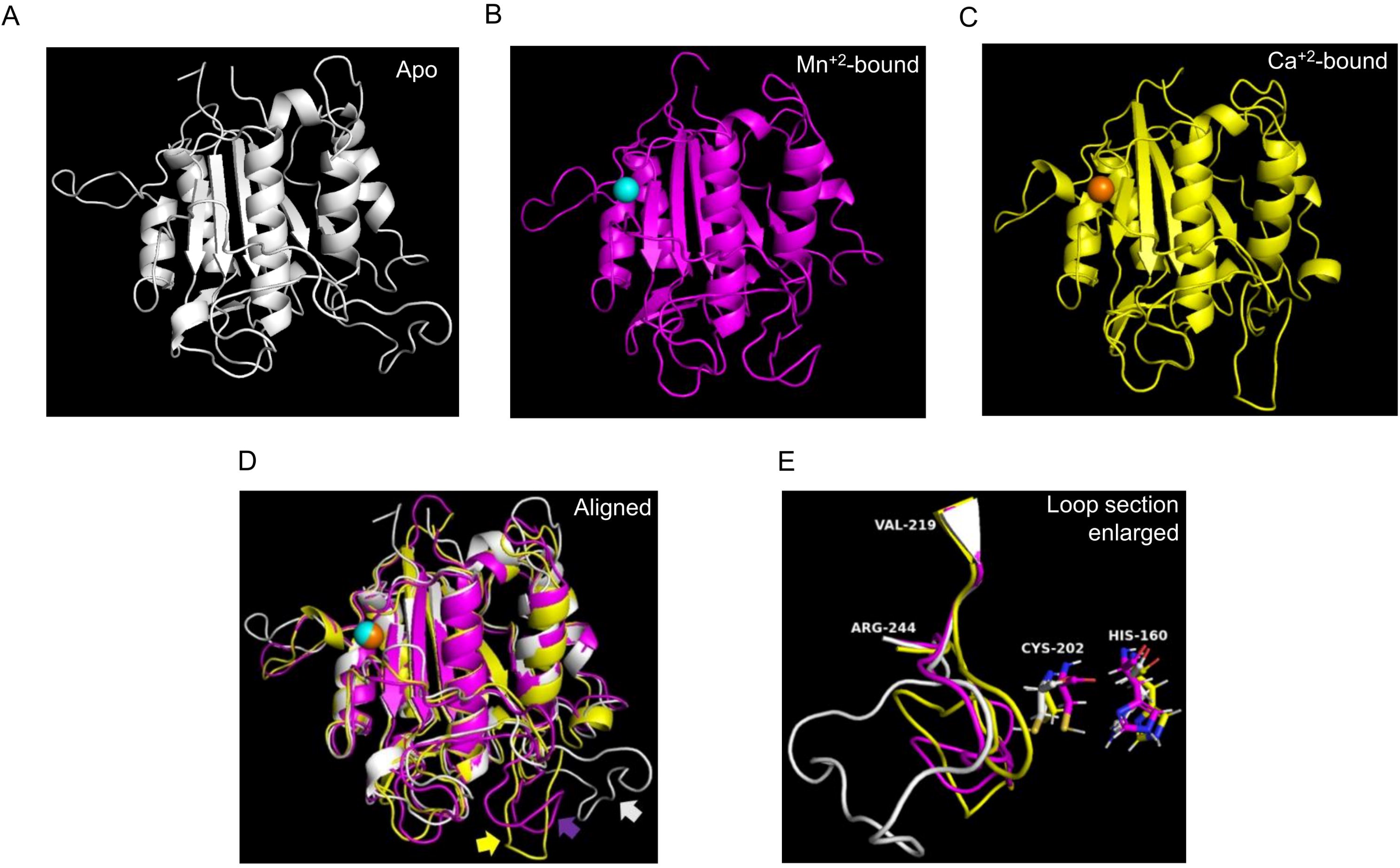
Snapshots of the metal-bound models of Gpi8_(36-303)_. The 100 ns frame of **(A)** apo Gpi8_(36-303)_, **(B)** Mn^+2^-bound Gpi8_(36-303)_ and **(C)** Ca^+2^-bound Gpi8_(36-303)_ models as visualized in PyMol. **(D)** Overlapped images of all the three 100 ns structures to visualize structural deviations. The apo form is represented in white, Mn^+2^-bound form in magenta and Ca^+2^-bound form in yellow. Mn^+2^ is represented in cyan and Ca^+2^ is represented in orange. Different coloured arrows (white for apo, magenta for Mn^+2^-bound and yellow for Ca^+2^-bound model) was added to incorporate the loop fluctuations near the active site in 100 ns structures. **(E)** Enlarged view of the loop region 219-244 and the catalytic dyad residues (Cys202 and His160) to visualize loop fluctuations.

We were particularly interested in a long flexible loop region between 219-244 residues (VGSSEIEESSYSHHSDMDIGVAVIDR) present in the vicinity of the catalytic pocket which is rich in amino acids capable of forming hydrogen bond networks (Figure 8E). The Ca^+2^-bound model appears to be the most compact and very minor structural fluctuations are observed throughout the simulation even within the loop region. The catalytic pocket is well assembled and no major structural disorganization is observed. In the Mn^+2^-bound model, some fluctuations are observed throughout the simulation, particularly in the 219-244 loop, which tends to swing back and forth in the vicinity of Cys202, without covering the catalytic pocket and without causing any global changes in the overall structure of the protein. In the apo form, perhaps due to its less compact overall structure, this loop tends to move away from the catalytic site (Figure 8E) and open-up by the end of the run, indicating loop disassembly.

A nine-residue peptide, TIDTNENGS (including the ω+1 and ω+2 residues, C-terminal to the ω site), was docked into the model obtained at the end of the 100 ns molecular simulations run. After energy minimization, molecular simulation runs of 30 ns were further carried out. As can be seen, from the models obtained, the flexible loop of 219-244 residues could participate in substrate binding and help align the peptide with respect to the catalytic dyad (Figure 9A, Figure 9B). Further, we see that the substrate arranges itself in a different orientation in each of the three cases due to the different orientations of the loop residues within these structures. In the apo form, Arg56, Ser222, Glu226, Ser230, and His231 are the major residues involved in H-bonding with Thr4, Asn5 and Asn7 of the peptide substrate to position it within the catalytic pocket (Figure 9A(i), Figure 9B(i)). In the Mn^+2^-bound form (Figure 9A(ii), Figure 9B(ii)), Asn54, His57, Gly161, Asn163, Thr201, Glu225, Ser227 and Ser228, interact with Thr4, Asn5, Glu6, Asn7 and Ser9 of the peptide substrate to position it within the catalytic pocket. In addition, the catalytic residues, His160 (via its imidazole ring) and Cys202 (via its amide backbone) are seen to be interacting with Asn5 of the substrate via H-bonding. This probably contributes to correctly orienting the catalytic residues with respect to the scissile bond of the substrate. In the Ca^+2^ bound form (Figure 9A(iii), Figure 9B(iii)), the orientation of the peptide substrate is different from that in the Mn^+2^-bound form, resulting in a larger number of H-bonding interactions. The most prominent ones among these are by Arg56, His160, Ser230, Ser233, Gly238 and Ala240, which form H-bonds with Asp3, Thr4, Asn5, Asn7, and Gly8 of the peptide substrate as well as with both the ω+1 and ω+2 residues. There are also interactions that are much shorter lived in Ca^+2^-bound form in comparison to the other two forms perhaps due to its compact nature. Heat maps corresponding to the H-bonding interactions are shown in Figure 9C.

**Figure 9.**
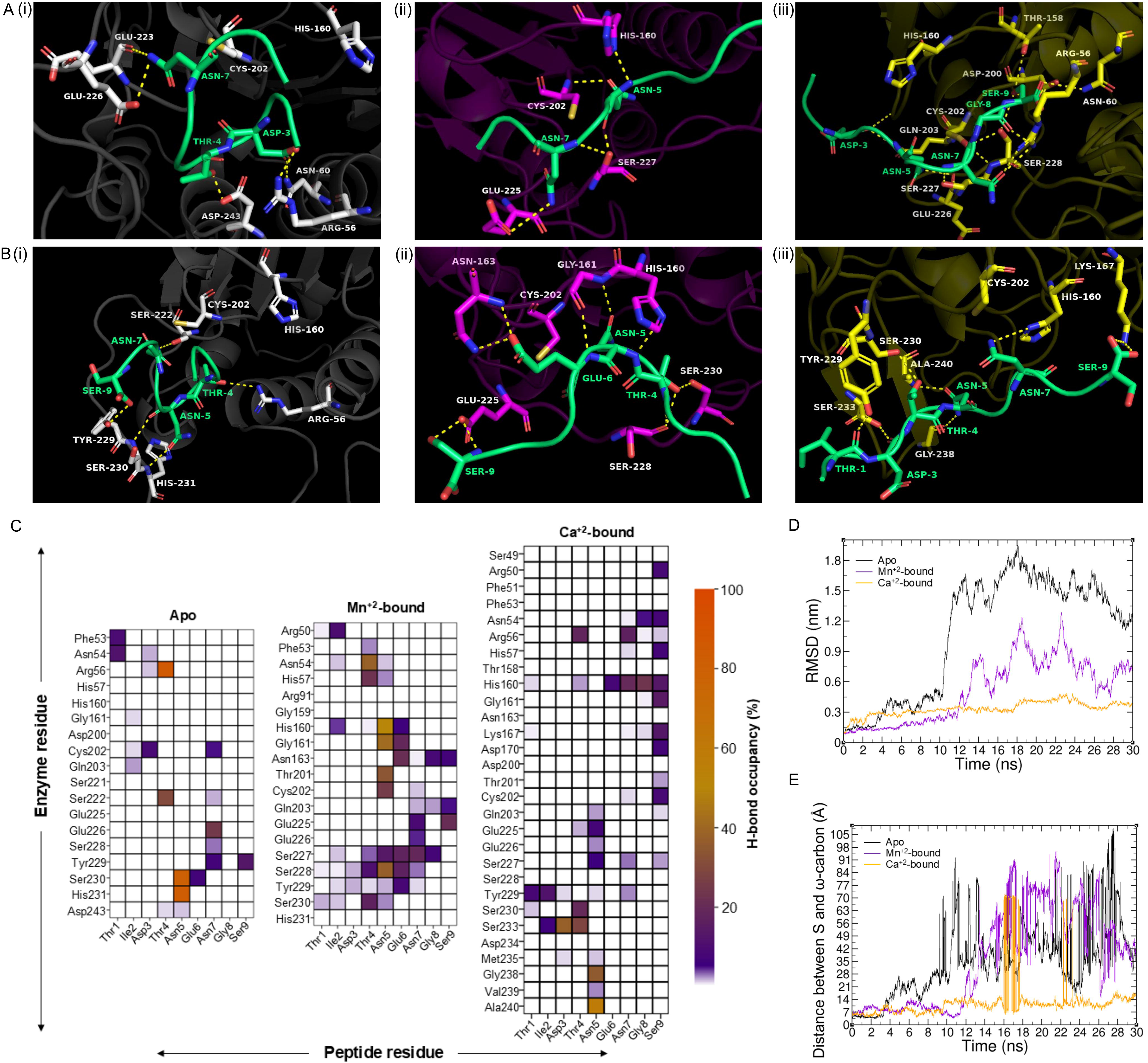
Snapshots of the peptide-docked models of Gpi8_(36-303)_. **(A)** Snapshot of binding site in docked model of **(i)** apoGpi8_(36-303)_, **(ii)** Mn^+2^-bound Gpi8_(36-303)_ and **(iii)** Ca^+2^-bound Gpi8_(36-303)_ models used for molecular simulation as obtained from HPEPDOCK protein-peptide docking server. **(B)** Snapshot of the binding site in those representative models of **(i)** apoGpi8_(36-303)_, **(ii)** Mn^+2^-bound Gpi8_(36-303)_ and **(iii)** Ca^+2^-bound Gpi8_(36-303)_ that show maximum number of hydrogen bond interactions. **(C)** Heatmaps for each model, generated by using data analysis and visualization tool from Python library (NumPy and Matplotlib), show possible hydrogen bond interactions of residues in the Gpi8_(36-303)_ substrate binding pocket with residues of the peptide (in percentage) until the time point where the peptide remains close to the catalytic Cys202. Interacting partners were identified using MDAnalysis. **(D)** Root Mean Square Deviation (RMSD) analysis shows higher structural deviations in peptide-docked apoGpi8_(36-303)_ as compared to the metal bound forms of Gpi8_(36-303)_. **(E)** Distance overlay plot show that the distance between −SH group of Cys202 and ω-carbon of scissile peptide bond of the substrate changes across the simulation run. The plots were generated by XMgrace.

The RMSD data confirm that structural fluctuations were higher in the apo form in comparison to metal bound forms (Figure 9D). For the apo form, these fluctuations begin immediately after 3 ns and increase further beyond 10 ns. This might indicate that the peptide is weakly bound to the catalytic site and could leave the pocket earlier than expected. In the Mn^+2^-bound form, RMSD was stable till 12 ns. The increased RMSD beyond this time point could imply that the peptide leaves the pocket after 12 ns. For the Ca^+2^-bound form, there is slight increase in the RMSD value in the initial phase relative to the other two models but it remains stable till the end of the run, suggesting that the peptide remains bound until the end of the run.

#### 3.5.5. Peptide substrate remains close to Cys202 in the metal bound forms

Next, we examined the distance between the catalytic Cys202 and the scissile bond of the substrate (between Asn7-Gly8). In the apo form, the peptide substrate remains close to the catalytic residue (Cys202) for only up to 3 ns and then begins to drift away from it (Figure 9E). As the peptide is close to the pocket only in the initial stages where pose repositioning might be happening, it confirms the possibility that the peptide binding is weak in the apoGpi8. On the other hand, both the metal-bound forms tend to show stable interaction till about 11 ns and 9.5 ns, respectively, for Mn^+2^- and Ca^+2^-bound Gpi8. It is also observed that in the metal-bound forms, although the distance fluctuates within the time frame, there exist conformations even at later time points where the peptide tends to get close to the catalytic Cys202 rather than moving away from it. This trend is seen in both the cases of Mn^+2^-bound and Ca^+2^-bound form, where early in the time frame the peptide starts to move away from the catalytic site but later shows a tendency to reoccupy the catalytic site. This is only possible when other interactions in the vicinity of the catalytic site are efficient enough to favour this.

From the docking poses, it was observed that the minimum distance between the thiol group of Cys202 and ω site residue/scissile peptide bond of substrate is about 3.56 Å at 10.64 ns in the case of Mn^+2^-bound Gpi8 and 3.95 Å at 6.70 ns for the Ca^+2^-bound form. In order to examine whether the average distance between Cys202 and scissile peptide bond is different in the two metal-bound forms, ten frames with the lowest distances for peptide bond cleavage were extracted and averaged. Thus, we estimated that the average minimum distance between Cys202 and the scissile bond was 3.8 ± 0.1 Å for the Mn^+2^-bound and 4.2 ± 0.1 Å for the Ca^+2^-bound forms. Perhaps the lower ionic radius of Mn^+2^ over Ca^+2^, could explain these conformational differences and the reason why the Mn^+2^-bound form is catalytically more efficient.

## 4. Conclusion

*C. albicans* is an opportunistic pathogenic fungus which can exist as a commensal in the human body but can produce infections in immunocompromised individuals that can be fatal. For host-recognition and pathogenesis, it requires a diverse array of virulence factors, many of which are expressed as GPI-anchored proteins. Production of GPI anchored proteins requires the pre-formed glycolipid anchor to be attached to the C-terminal end of proteins. The GPIT performs this final role in a multi-step sequential pathway. This crucial step is required for the GPI-anchored virulence factors to be transported via the secretory pathway and be displayed at its cell surface.

The GPIT needs to proteolytically cleave a C-terminal signal sequence of proproteins and amide-link the newly created carboxylate end to the GPI precursor. While the sequence of the SS is not conserved, its overall nature is reasonably well conserved across eukaryotes. Thus, we know that the site of cleavage (ω site) and its flanking residues (ω-1 and ω+1) are amino acids with relatively small side-chains (eg. Gly, Ala, Asn, Asp, Cys, Ser), which are followed immediately at the C-terminus by 5-10 polar residues and then by a 15-20 residue hydrophobic tail [1,30–32]. Most prediction programs available online exploit these features of the SS [27–29]. The molecular details of this preference are not well understood. This study attempts to shed some light on these aspects using *C. albicans* GPIT. Specifically, we examined why the size of the residue at the ω site and the length of the peptide segment containing it matters. We also explain why, despite not being required for catalytic activity *per se*, the choice of the divalent cation matters for the endopeptidase activity.

In order to do this, we employed a modified cell free assay protocol from the one we reported previously [15]. Instead of precipitating out microsomes from PMF with CaCl_2_, we used the PMF samples directly. This ensured that our samples did not have excess Ca^+2^ ions to begin with. Using freshly generated PMF samples and TN7-AMC in the assay, we show that the endopeptidase activity is Gpi8-specific, and shows comparable trends as what was observed with the microsomal assays.

We then show that EDTA treatment of the PMF reduces the endopeptidase activity substantially, but doesn’t irreversibly inactivate it. The addition of Mn^+2^ after EDTA treatment results in higher activity than observed in freshly prepared PMF, suggesting that the endogenously bound cofactor may be different from Mn^+2^ or that the process of preparing the PMF causes at least some of the GPIT complexes to lose their bound divalent cations initially and which are restored upon reconstitution with the divalent cation.

Steady state analysis using TN7-AMC suggests that Mn^+2^ reduces K_M_ without significantly altering the V_max_. In other words, the divalent cation does not play a direct role in altering catalytic rates even though it enables better substrate binding. This is substantiated by the fact that the divalent cation did not alter the optimum pH of the enzyme, which would be expected if it played a role in generating the nucleophile.

Could it be that the divalent cation enables coordination of the peptide substrate to residues within the catalytic pocket? We saw no substantial effect of Mn^+2^ on the fluorescence spectra of either the substrate or the product, to suggest that it might be interacting with them in any way. The structure of mammalian PIG-K has a bound Ca^+2^ ion at a site distant from the active site [11]. Molecular dynamic simulations on a model for soluble domain of Gpi8 (Gpi8_(36-303)_) generated by AlphaFold Server 3, which compared well with the PIG-K structure, suggested that the role of the divalent cation was to stabilize the native structure without significantly altering the overall steric fit or the hydrophobic packing within the folded protein core.

Peptide-docking and molecular dynamic simulations also suggested that the conformation of a flexible loop between residues 219-244 was significantly different in the metal-bound forms versus the apo form. In the former, this loop was positioned such that it not only interacted with the peptide substrate and oriented it within the active site, but it also enabled the substrate to reoccupy the catalytic pocket when drifting away from it. Such an enabling interaction for the substrate was absent in apoGpi8. Based on these results, we propose that the role of the divalent cation is two-fold: 1) to improve substrate binding by stabilizing the conformation of the enzyme and, 2) to enable a conformation that positions loop 219-244 in the vicinity of the binding pocket for stabilizing the bound substrate via H-bonding. Of the two divalent cations tested, Mn^+2^ bound to Gpi8 created a slightly less rigid structure than did Ca^+2^. Its smaller ionic radius also appeared to enable an active site geometry in which the scissile bond of the peptide substrate can approach closer to the catalytic thiol of Cys202 than in Ca^+2^-bound form of the protein. This probably explains why Mn^+2^ is the preferred cation for stimulating endopeptidase activity in our cell free assays.

Next, we examined the dependence of peptide substrate length on the endopeptidase activity of *C. albicans* GPIT. AMC-tagged 15-mer, 9-mer, 7-mer and 4-mer peptides were custom-synthesized for the analysis based on the Hyr1 GPI attachment SS, such that they were all elaborations or deletions of TN7. We found that peptides that were 7-mer to 9-mer in length were better substrates than either the shorter or the longer substrates tested. This clearly indicated that the length of the substrate fitting into the catalytic pocket mattered. A GPIT complex of *S. cerevisiae* consisting only of soluble truncated mutants of Gpi8, Gaa1, and Gpi16 was also shown to most optimally process an 8-mer substrate over a 4-mer or a 19-mer [14]. This is not surprising, since cysteine proteases are known to have extended substrate binding sites that stabilize and correctly position the substrate within the pocket [33]. Indeed, using a 9-mer peptide in our molecular dynamic simulations also demonstrated extensive interactions of the N-terminal residues Asp3, Thr4, and Asn5 (residues P4, P3, P2) of the peptide substrate with the catalytic pocket in the Mn^+2^-bound Gpi8 such that Asn7 and the scissile bond (between P1-P1’) C-terminal to it were positioned close to Cys202. This would also explain why GPIT in the *C. albicans* PMF samples had better binding affinity for VN9-AMC and TN7-AMC relative to TN4-AMC. The 4-mer peptide probably makes fewer stabilizing contacts with the extended binding site and with the 219-244 flexible loop, which explains why, treatment of the PMF with EDTA did not substantially affect K_M_ for it. However, this also probably made it that much easier for the cleaved peptide product to leave the catalytic pocket, resulting in higher V_max_ values. The addition of divalent cations, Mn^+2^ or Ca^+2^, to this system restores rigidity of the protein structure and reduces catalytic rates.

What explains the weaker affinity of the enzyme for a 15-mer peptide? Why doesn’t it bind the catalytic pocket with similar (if not better) affinity as the 9-mer, given that theoretically it too could have established similar contacts within the catalytic pocket? One hypothesis would be that this longer peptide spontaneously folds in aqueous solution, adopting a conformation that does not allow it to efficiently interact with the catalytic pocket.

Should the size of the residue at the site of cleavage alter the kinetics of endopeptidase activity? A 7-mer peptide with the ω site residue changed from Asn (as in TN7) to Pro (as in TP7) bound with a much weaker K_M_ than a 7-mer or 9-mer, suggesting that a bulky residue at the site of cleavage probably experiences steric hindrance in fitting into the catalytic pocket. A similar preference for amino acids with smaller side chains is well established in the case of GPIT from both *S. cerevisiae* and humans [30–32], and appears to be one of the underlying mechanisms by which specificity is built into this system.

One other issue that puzzled us was the low overall activity of the enzyme in the cell free assays. The simulation studies provide some clues as to what might be occurring. The active site of the enzyme appears to be a solvent-exposed hollow pocket allowing short peptide substrates to diffuse away easily. Thus, the number of productive interactions at the active site are probably low. In the case of the cellular system, the SS is normally 20-30 amino acids long and is expected to be inserted in or associated with the ER lumenal membrane via its hydrophobic tail. Cryo-EM structures of mammalian and *S. cerevisiae* GPIT suggest that the SS peptide substrate is bound in a narrow channel with stabilizing interactions established by Gpi16 (PIG-T) and Gpi17 (PIG-S) homologs [11,34]. In the absence of the hydrophobic tail the peptide substrates used in this study are probably poorly inserted into the ER membrane fragments of the PMF and are perhaps not guided efficiently into the pocket.

Nevertheless, our study suggests that the cell free system can correctly reflect some of the characteristic features expected of the GPIT, its co-factor specificity, preference for an amino acid with a small side chain at the ω-site and its requirement for peptides of optimal length for effective binding within the active site.

## Acknowledgements

Parts of this work were funded by grants to S.S.K. from Department of Biotechnology (DBT) India BT/PR53604/BMS/85/165/2024, BT/PR51192/MED/29/1662/2023 (as co-PI) and the SERB-POWER Fellowship (SPF/2021/000097). IC was supported by PMRF fellowship and funding. SS received a salary from a DBT-sponsored project (BT/PR39195/BRB/10/1889/ 2020) to S.S.K. Fluorescence spectroscopy was performed at the Central Instrumentation Facility (CIF), School of Life Sciences (SLS), JNU. Facilities and research at SLS have been supported by grants from UGC-CAS, UGC-RNW, DBT-BUILDER, and DST-FIST.

## CRediT authorship contribution statement

**Isaac Cherian**- Generated the data and analyzed the data shown in Table III, Figures 1-5, Figure 6B, Figure 6C, Figures 7-8, Figure 9 (A-C & E), and all the supplementary figures.

**Shailja Shefali**- Standardized the enzyme assay protocols and contributed to the background studies.

**Dip Shikha Maurya**- Worked on characterizing the enzymatic assay using PMF for her Masters dissertation and contributed to the background studies.

**Faraz Ahmed**- Performed the molecular dynamic simulations (Figure 6A, 6D, 6E and Figure 9D).

**Sneha Sudha Komath**- Conceptualised and supervised the project, wrote the overall framework of the manuscript, and brought in the funds required.

All authors have read, edited/ commented and agreed with the final version of the manuscript.

## Supporting Information

**Figure S1.** Fluorescence emission of the peptide substrates are unaffected by EDTA or by the addition of 5 mM divalent cations.

**Figure S2.** Fluorescence emission of AMC is unaffected by EDTA or by the addition of 5 mM divalent cations.

**Figure S3.** AMC standard curve for determining concentration of product formed in the endopeptidase assay.

**Figure S4.** The structure of Gpi8_(36-303)_ compares well with the cryo-EM structure of PIG-K.

