## Supplementary File for "Characterizing the endopeptidase activity of *Candida albicans* Gpi8, a crucial subunit of the GPI transamidase"

### Supplementary Figures

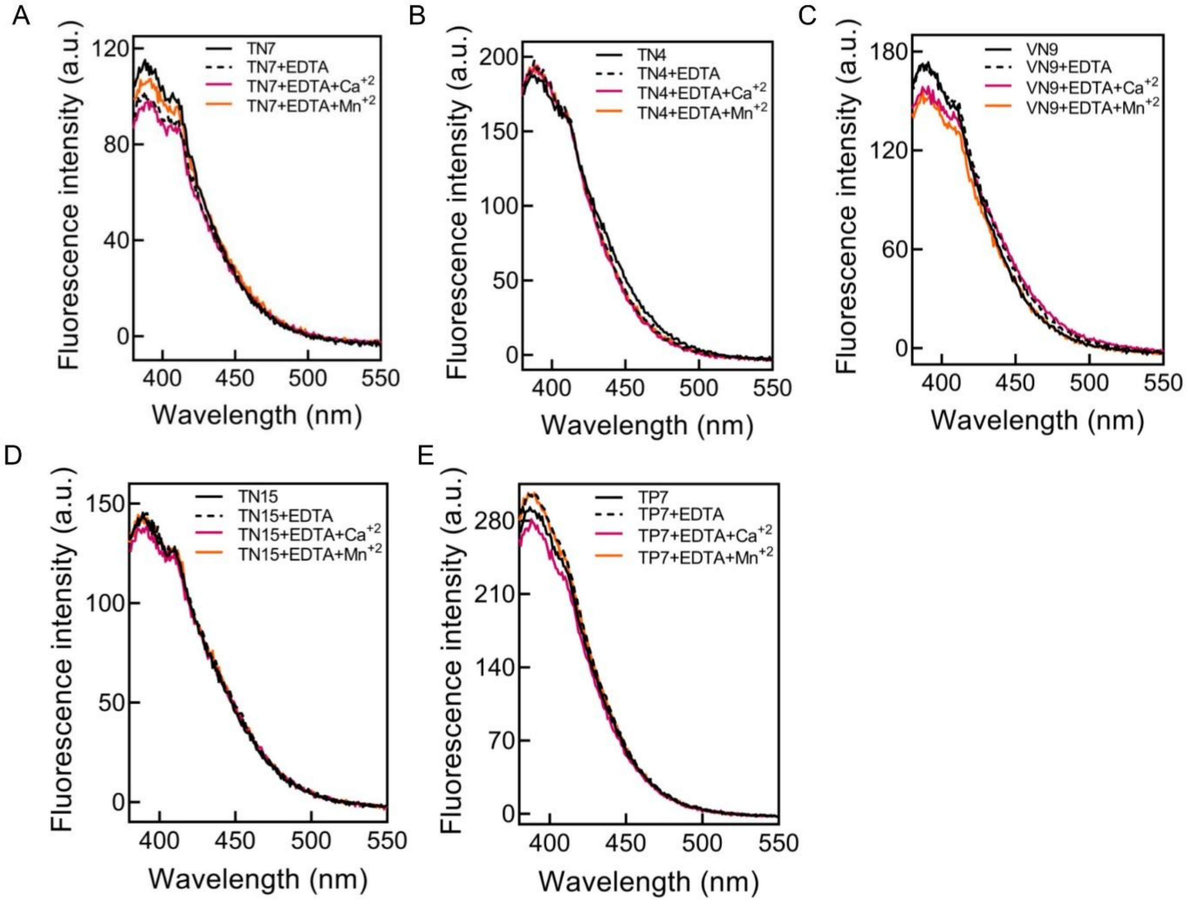


**Figure S1. Fluorescence emission of the peptide substrates is unaffected by EDTA or by the addition of 5 mM divalent cations.** Spectra showing stability of **(A)** TN7-AMC **(B)** TN4-AMC **(C)** VN9-AMC **(D)** TN15-AMC and **(E)** TP7-AMC in presence of EDTA, EDTA+$Mn^{+2}$ and EDTA + $Ca^{+2}$. The samples were incubated for 6 h at 37 °C in 100 mM Tris-HCl (pH 7.2). and fluorescence emission spectra were recorded between 380-550 nm after excitation at 360 nm. Slit widths of 3 nm were used for excitation and emission. Note: a.u. represents arbitrary units.

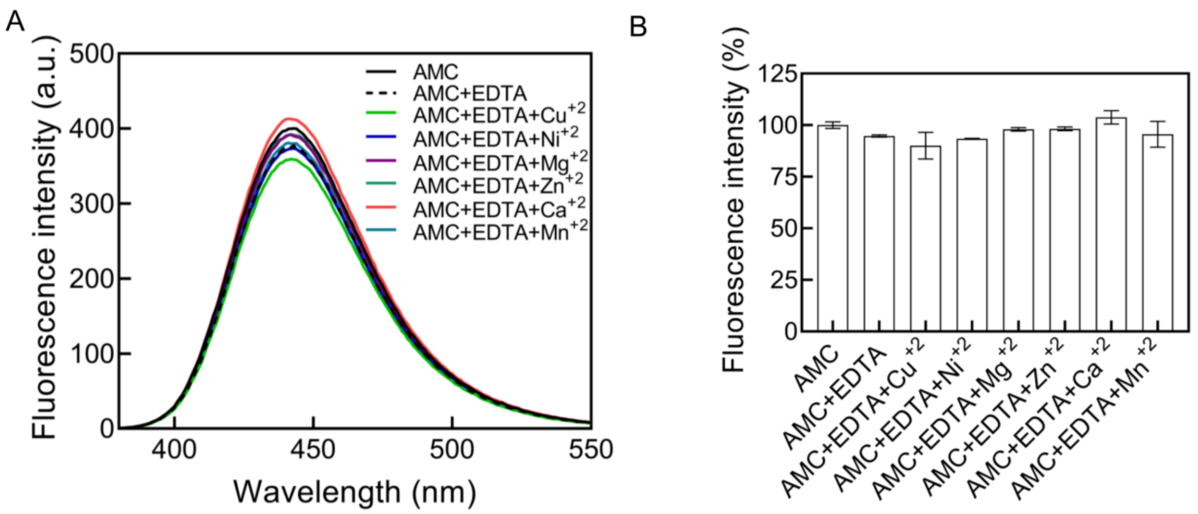


**Figure S2. Fluorescence emission of AMC is unaffected by EDTA or by the addition of 5 mM divalent cations.** AMC was incubated for 6 h at 37 °C in 100 mM Tris-HCl (pH 7.2) in the absence or presence of EDTA, or EDTA + 5 mM solutions of divalent cation salts ($CuCl_2$, $ZnCl_2$, $MgCl_2$, $NiCl_2$, $CaCl_2$ or $MnCl_2$). Fluorescence emission spectra were recorded between 380-550 nm after excitation at 360 nm. Slit widths of 3 nm were used for excitation and emission. **(A)** Representative spectra of the samples as shown in the figure. **(B)** Bar graphs showing relative fluorescence emission intensity at 440 nm in AMC+EDTA and AMC+EDTA+divalent cations relative to that of AMC alone in the same buffer (as percentages). None of the data showed statistically significant differences. Note: a.u.; arbitrary units.

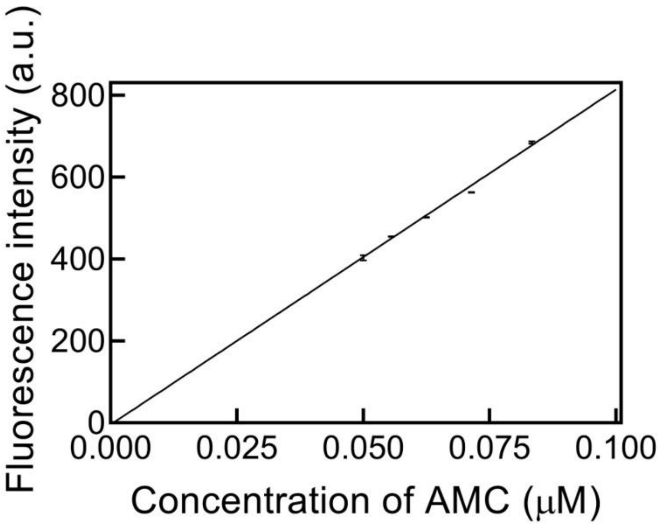


**Figure S3. AMC standard curve for determining concentration of product formed in the endopeptidase assay.** Different concentrations of AMC were taken in 100 mM Tris-HCl (pH 7.2). Fluorescence emission spectra were recorded between 380-550 nm after excitation at 360 nm. Slit widths of 3 nm were used for excitation and emission. From the spectra, the fluorescence emission intensity at 440 nm was determined and used to plot a graph of Fluorescence intensity v/s concentration of AMC. This was used to determine the concentrations of our experimental samples based on their fluorescence emission at 440 nm. Since fluorescence is measured in arbitrary units, for each experiment a standard AMC curve needed to obtained in parallel. The data shown in this plot is representative of one such standard curve done in duplicates. Note: a.u.; arbitrary units.

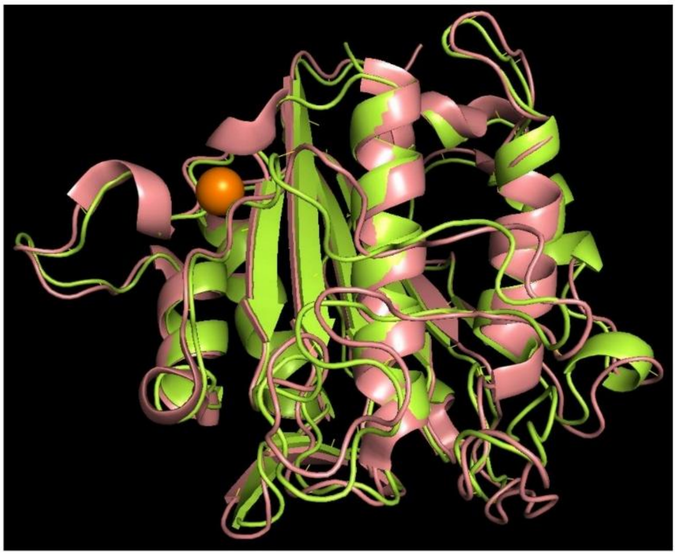

**Figure S4. The structure of Gpi8$_{(36-303)}$ compares well with the cryo-EM structure of PIG-K.** Alignment of human PIG-K obtained from cryo-EM structure of GPI transamidase (PDB ID- 7WLD) with CaGpi8$_{36-303}$ generated by AlphaFold server show that they are structurally well aligned (RMSD of 0.757). The protein structure for PIG-K is represented in salmon red, that for CaGpi8$_{36-303}$ in limon, and $Ca^{+2}$ in orange.
